# Translation rescue by targeting Ppp1r15a upstream open reading frame *in vivo*

**DOI:** 10.1101/2021.12.11.472232

**Authors:** Ashley Kidwell, Shiv Pratap Singh Yadav, Bernhard Maier, Amy Zollman, Kevin Ni, Arvin Halim, Danielle Janosevic, Jered Myslinski, Farooq Syed, Lifan Zeng, Alain Bopda Waffo, Kimihiko Banno, Xiaoling Xuei, Emma H. Doud, Pierre C. Dagher, Takashi Hato

## Abstract

The eIF2 initiation complex is central to maintaining a functional translation machinery. Extreme stress such as life-threatening sepsis exposes vulnerabilities in this tightly regulated system, resulting in an imbalance between the opposing actions of kinases and phosphatases on the main regulatory subunit eIF2α. Here, we report that translation shutdown is a hallmark of established sepsis-induced kidney injury brought about by excessive eIF2α phosphorylation and sustained by blunted expression of the counterregulatory phosphatase subunit Ppp1r15a. We determined that the blunted Ppp1r15a expression persists because of the presence of an upstream open reading frame (uORF). Overcoming this barrier with genetic approaches enabled the derepression of Ppp1r15a, salvaged translation and improved kidney function in an endotoxemia model. We also found that the loss of this uORF has broad effects on the composition and phosphorylation status of the immunopeptidome that extended beyond the eIF2α axis. Collectively, our findings define the breath and potency of the highly conserved Ppp1r15a uORF and provide a paradigm for the design of uORF-based translation rheostat strategies. The ability to accurately control the dynamics of translation during sepsis will open new paths for the development of therapies at codon level precision.

## Introduction

Sepsis-induced multi-organ failure is a major life-threatening condition with no effective therapy. The current paradigm views inflammation as the centerpiece of early sepsis pathology^1-3^. However, this paradigm falls short in connecting inflammation to severe organ failure that is frequently observed in the late phase of sepsis. A crucial molecular event underpinning this transition is phosphorylation of the main translation regulator eIF2α^4,5^. Phosphorylated eIF2α causes nearly all translation initiation processes to be stalled because phosphorylated eIF2α inhibits GTP/GDP exchange required for recycling of eIF2 between successive rounds of initiation. Transient inhibition of protein synthesis could be cytoprotective as it attenuates energy consumption and upregulates the adaptive response to stress (the integrated stress response). However, persistent inhibition of global protein synthesis is clearly detrimental^6,7^. The optimal dynamics of eIF2α phosphorylation required for successful recovery from sepsis-induced organ failure is not known.

In sepsis, eIF2α phosphorylation is primarily mediated by EIF2AK2/PKR (Protein Kinase, Interferon-Inducible Double Stranded RNA Dependent; **Suppl Fig. 1A**)^8,9^. Dephosphorylation of eIF2α is catalyzed by eIF2α holophosphatases that are composed of ubiquitous protein phosphatase 1 (catalytic subunit) bound to one of two regulatory subunits (Ppp1r15a, also known as GADD34, and Ppp1r15b)^10,11^. Of the two regulatory subunits, only Ppp1r15a is induced by stress, making it a prime therapeutic candidate for this strategic rescue of translation in late phase sepsis. Here, we report that Ppp1r15a is translationally repressed during late phase sepsis due to the existence of an upstream open reading frame (uORF) in the 5’ leader sequence of Ppp1r15a. Using genetic and oligonucleotide approaches, we characterize the significance of the uORF in blunting Ppp1r15a expression and causing sustained translation shutdown. Our work provides insights into how a single uORF can deeply impact the pathophysiology of sepsis and serves as a distinct layer of molecular regulation in health and disease.

## Results

### Ppp1r15a uORF is a translational repressor of its CDS

Progression and recovery of sepsis-induced kidney injury correlate with changes in global translation^12^. The early phase of kidney injury is characterized by an increase in overall translation (**Fig. 1A**). Tissue-derived proinflammatory cytokines and chemokines are rapidly produced during this time^12-14^. The resultant stress responses converge on the phosphorylation of the main translation regulator eIF2α (**Fig. 1B**), leading to translation shutdown at later timepoints in sepsis (12 – 18 hrs after LPS/endotoxemia and CLP/cecal ligation and puncture in mice)^12^. This translation shutdown is triggered by activation of antiviral responses that occur irrespective of the specific initiating pathogen^8,15^. Importantly, pharmacologic reversal of this late translation shutdown accelerates renal recovery, suggesting that defective protein synthesis is a driver of sustained tissue damage rather than a byproduct^12^. Using our published Ribo-seq data, here we searched for a molecular target that could be contributing to this sustained translation shutdown. We identified that the expression of Ppp1r15a (Protein Phosphatase 1 Regulatory Subunit 15A) was repressed during endotoxemia (**Suppl Fig. 1B**). Ppp1r15a is a key counter regulatory molecule required for the reversal of eIF2α phosphorylation. Thus, blunted Ppp1r15a expression could explain delayed recovery from translation shutdown. Comparison of Ribo-seq, RNA-seq, and ATAC-seq data indicates that Ppp1r15a expression is translationally repressed in sepsis possibly because of the presence of an uORF in front of the canonical coding sequence (CDS; **Suppl Fig. 1B-D**).

**Figure 1.**
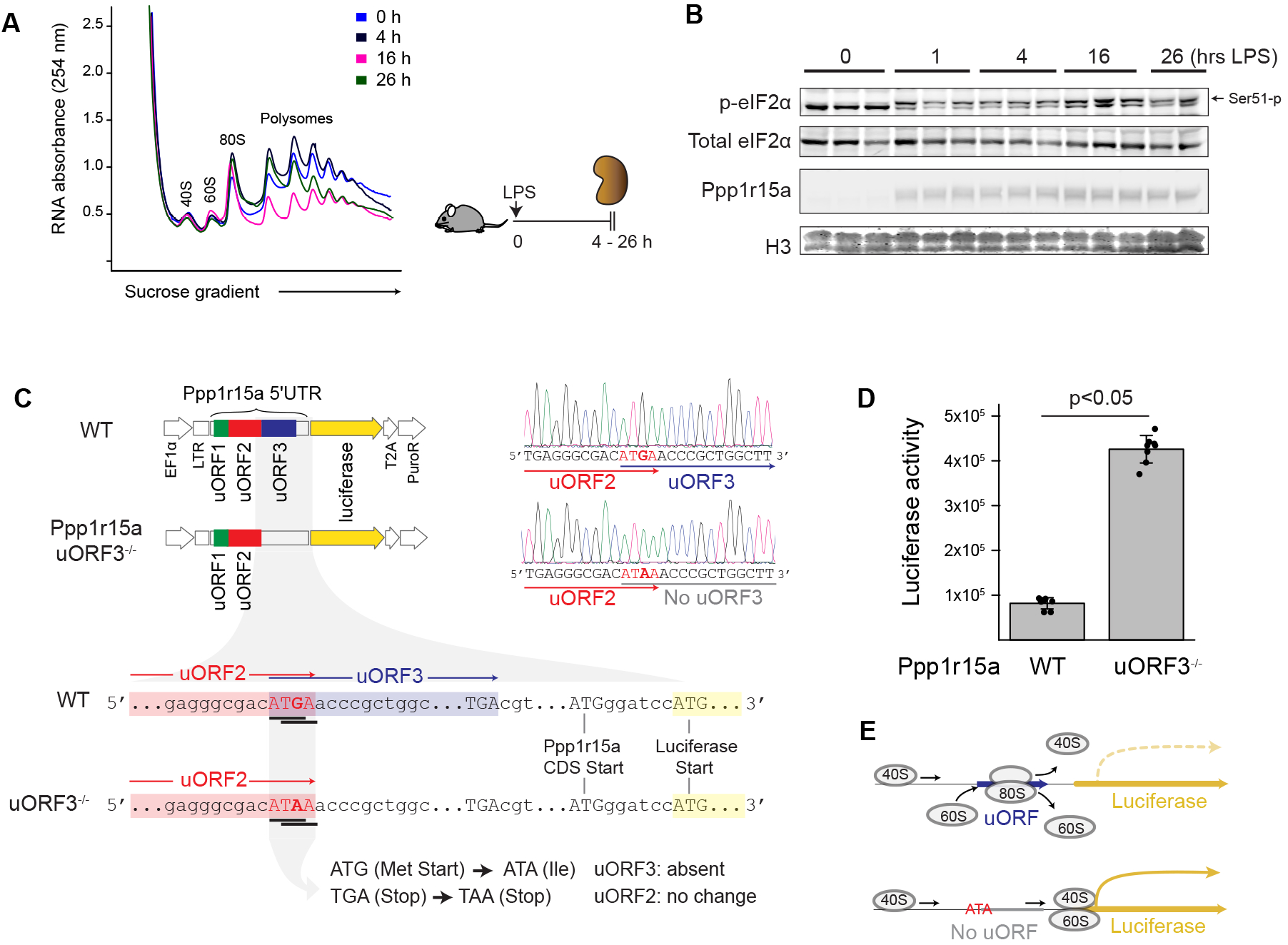
Ppp1r15a uORF inhibits translation of Ppp1r15a CDS. (**A**) Polyribosome profiling of kidney extracts from mice treated with LPS 5 mg/kg i.v. for indicated durations. Ribosomal subunit 40S, 60S, mono-ribosome (80S), and polyribosomes were separated using sucrose density gradient (10-50%). Polyribosome signals peaked at 4 hrs and reached a nadir at 16 hrs after endotoxin challenge. (**B**) Time-course analysis of eIF2α serine51 phosphorylation and Ppp1r15a by western blot is shown. H3, histone. (**C**) Reporter constructs used in this study and the Sanger sequencing chromatograms. Full-length mouse Ppp1r15a 5’UTR was fused to firefly luciferase in frame on the pCDH backbone vector. In the mutant construct, the 3^rd^ uORF (uORF3) was abolished by introducing a single nucleotide mutation in the start codon. (**D**) Luciferase activity levels are shown for cells transfected with Ppp1r15a 5’UTR WT and mutant plasmids. (**E**) Schematic of ribosome coverage over the constructs.

A wide range of uORFs have been implicated in cis-acting translational control of their main CDS^16-20^. Conversely, thousands of other uORFs are not characterized or do not have such regulatory function. For example, Ppp1r15b, the isoform of Ppp1r15a, exhibited ribosome-occupying uORF similar to Ppp1r15a (**Suppl Fig. 1B**). However, despite the robust ribosome engagement with the uORF, translation efficiency of Ppp1r15b CDS remained unchanged throughout the course of sepsis. These findings support the notion that uORF-mediated translational control is gene and context dependent^21,22^. Therefore, to determine the functionality of Ppp1r15a uORF, we designed plasmid constructs consisting of intact (wild-type) and mutant 5’ leader sequences of Ppp1r15a inserted between a promoter and the firefly luciferase reporter CDS in frame (**Fig. 1C**). We focused on the 3^rd^ uORF (uORF3, closest to the CDS start site) because of its prominent ribosome-occupancy and high sequence conservation across species (**Suppl Fig. 1E**, mouse uORF3 corresponds to human uORF2). In the mutant construct, uORF3 start codon was abolished by a single nucleotide mutation while maintaining the overlapping uORF2 stop codon intact. We found that elimination of the uORF3 start codon signal substantially increased the rate of downstream CDS translation (**Fig. 1D**), consistent with a model in which Ppp1r15a uORF serves as a barrier rather than an enhancer of its CDS translation^23,24^ (**Fig. 1E**). In summary, amidst sepsis-induced eIF2α phosphorylation and translation shutdown, counterregulatory Ppp1r15a remains suppressed due in part to the presence of its uORF.

### Length- and sequence-specific effects of Ppp1r15a uORF

To elucidate the role of endogenous uORF in the genome, we next generated PPP1R15A uORF mutant human cell lines using CRISPR-Cas9 with homology directed repair templates. These mutant cell lines include start codon mutant (effectively no uORF), short uORF (11 codons as opposed to 26 codons in wild-type), long uORF (99 codons), and substitution of difficult-to-translate polyprolines^25^ (wild-type PPP1R15A uORF harbors polyprolines at the C-terminus; **Fig. 2A-B, Suppl Fig. 2**). Similar to plasmid-based experiments, elimination of genomic uORF resulted in marked increases in the expression of PPP1R15A protein (No uORF, **Fig. 2C**). Despite the high PPP1R15A protein levels, overall translation profiles were comparable between no-uORF and wild-type at baseline (**Fig. 2D top**). This suggests that phosphorylated eIF2α, the target of PPP1R15A, is negligible in these cell lines under normal culture condition, or eIF2α is adequately dephosphorylated by constitutive expression of PPP1R15B. Truncation of uORF and elimination of polyproline resulted in higher levels of PPP1R15A protein expression as compared to wild-type, albeit less when compared to no-uORF. Conversely, long uORF resulted in minimal PPP1R15A protein expression (**Fig. 2C, Suppl Fig. 3A-C**). These findings indicate that both uORF length and polyproline content inversely impact the translation efficiency of downstream CDS.

**Figure 2.**
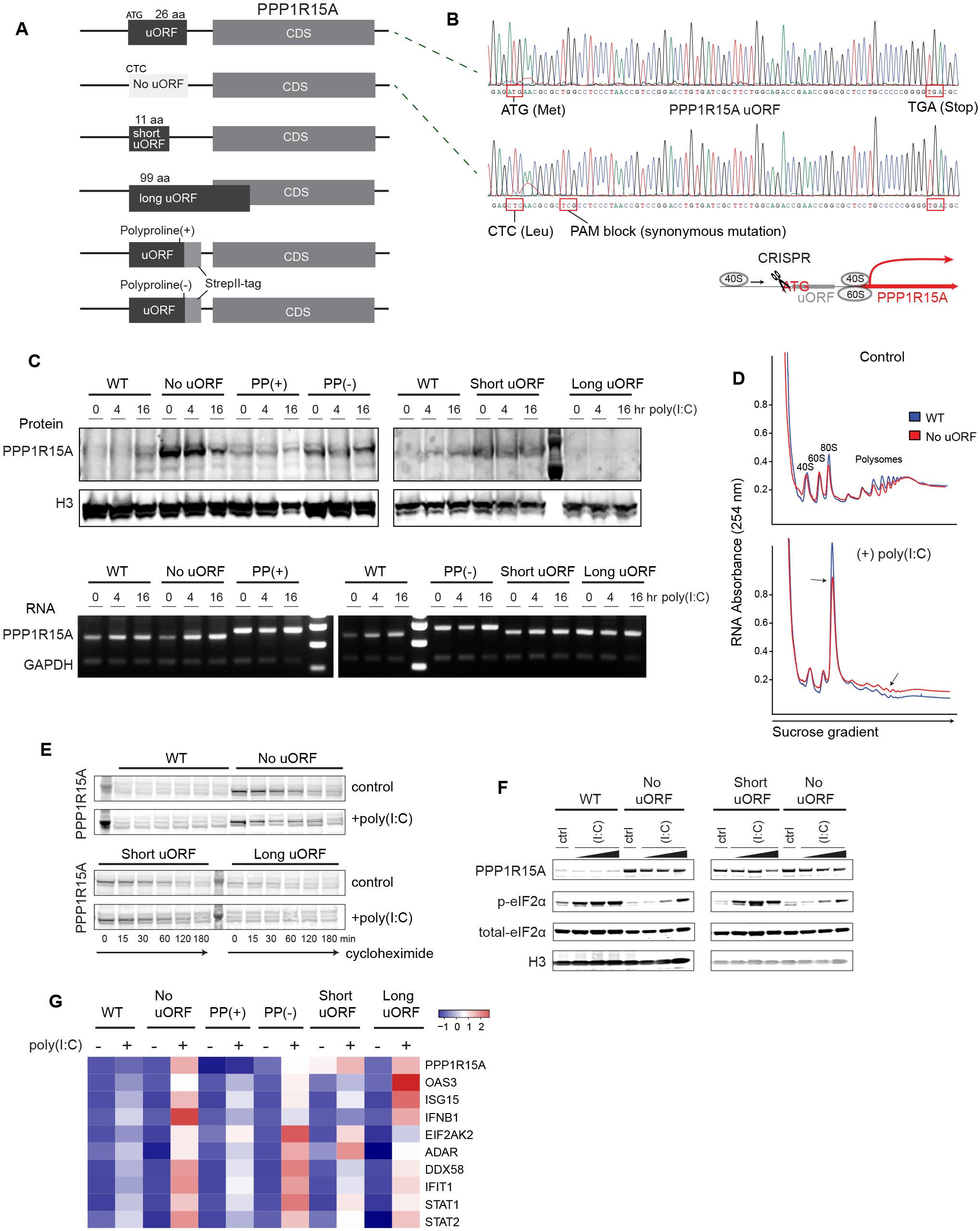
Modulation of Ppp1r15a uORF sequence differentially affects Ppp1r15a translation. (**A**) Schematic of mutant cell lines generated with CRISPR/Cas9. No-uORF cell line lacks uORF ATG start codon (instead CTC/Leucine). Short uORF harbors a truncated 11-mer uORF instead of 26 amino acids in wild-type. Long uORF encodes 99-mer uORF due to single nucleotide deletion and frame shift. Wild-type uORF ends with polyprolines (PP; proline-proline-glycine-stop). Polyproline(+) cell line retains this polyproline sequence followed by a linker and single Strep Tag II (proline-proline-glycine-linker-StrepTag-stop). Polyproline(-) cell line lacks this polyproline sequence, and instead glycine is inserted (glycine-glycine-glycine-linker-StrepTag-stop). (**B**) Sanger sequencing chromatograms for the indicated cell lines. Chromatograms for the rest of cell lines are shown in **Supplemental Figure 2**. (**C**) Western blot and PCR for PPP1R15A of indicated cell lines at baseline and after 1 μg/mL poly(I:C) transfection. The shift of PCR products in PP(+) and PP(-) is due to the StrepTag insert. (**D**) Polyribosome profiling of No-uORF cell line versus wild-type 16 hrs after poly(I:C) transfection. Arrows point to reduced monosome fraction and increased polysome fraction, indicating higher translation in No-uORF cell line. (**E**) Determination of PPP1R15A protein turnover. Nascent protein synthesis was inhibited with cycloheximide (250 μg/mL) for indicated durations with or without concomitant poly(I:C) treatment (1 μg/mL, 4 hrs). (**F**) Western blot for PPP1R15A and eIF2α under indicated conditions. Poly(I:C) concentrations used were 0.1, 0.5, and 1.5 μg/mL for 4 hrs. (**G**) RNA-seq analysis. Representative antiviral genes upregulated after 16 hrs of poly(I:C) treatment are shown (all within top 40 differentially expressed genes as shown in **Supplemental Figure 3A**).

**Figure 3.**
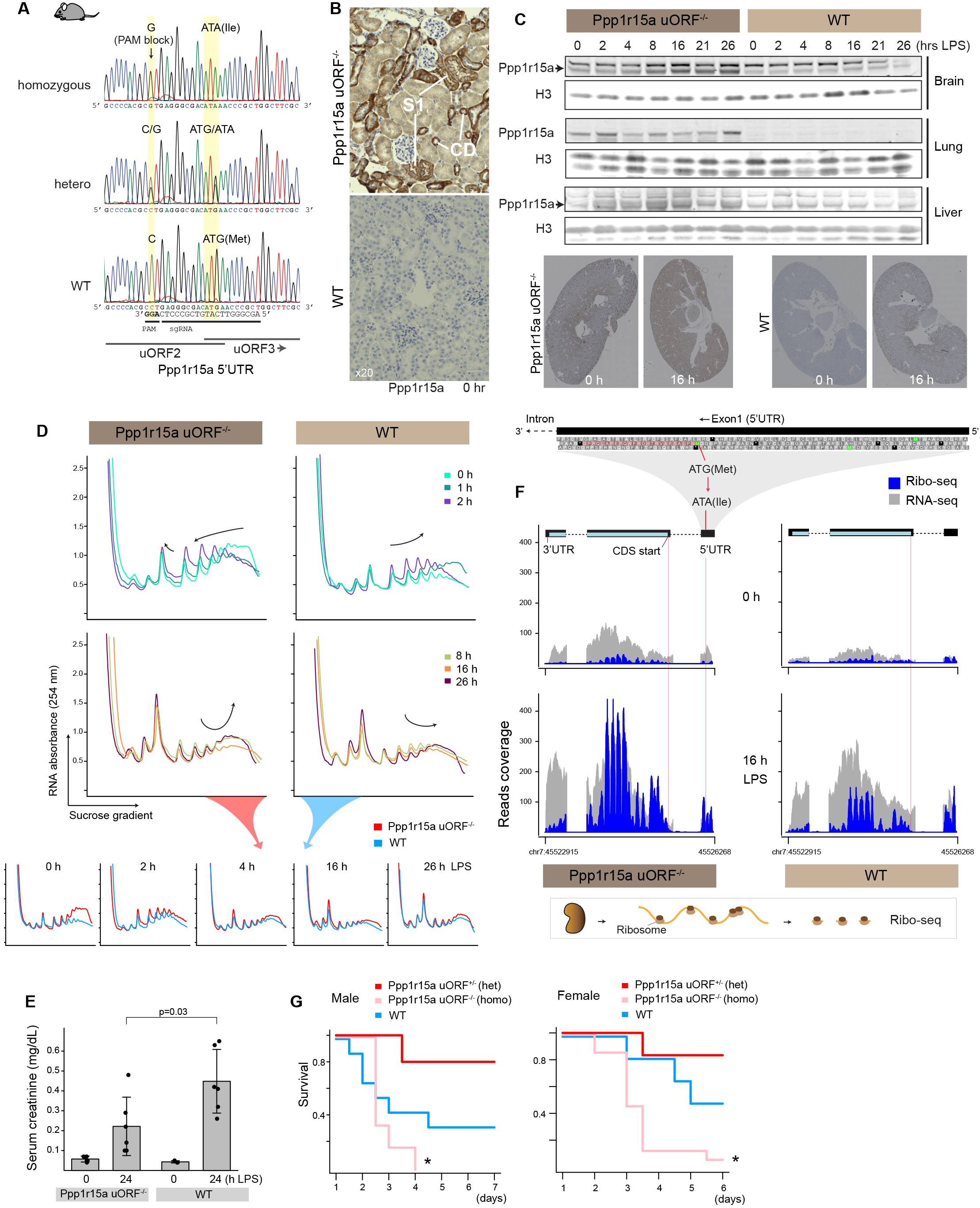
Phenotype characterization of Ppp1r15a uORF-deficient mice. (**A**) Sanger sequencing chromatograms of Ppp1r15a uORF mutant mice. (**B**) Representative immunohistochemistry staining of Ppp1r15a staining (brown) are shown for Ppp1r15a uORF mutant and wild-type mouse kidneys under indicated timepoints. Ppp1r15a signal is most pronounced in S1 proximal tubule and collecting duct (CD). (**C**) Western blot analysis of brain, lung, and liver in Ppp1r15a uORF mutant and wild-type under indicated timepoints after 5 mg/kg LPS tail vein injection. (**D**) Polyribosome profiling of kidney extracts from mice treated with LPS i.v. for indicated durations. For clarity, traces are overlayed within and between Ppp1r15a uORF mutant and wild-type mice for select timepoints. (**E**) Serum creatinine levels at baseline and 24 hrs after 5 mg/kg LPS i.v. for Ppp1r15a uORF mutant and wild-type male mice. (**F**) Ribo-seq analysis of Ppp1r15a under indicated conditions. RNA-seq reads are shown in gray and Ribo-seq reads in blue. (**G**) Survival curves after CLP are shown for homozygous, heterozygous Ppp1r15a uORF mutant and wild-type mice (n=7-9 per condition, *p <0.05 vs. heterozygous mice).

Exposure of cells to viral mimicry with poly(I:C) revealed a contrasting effect on PPP1R15A protein expression between wild-type and no-uORF cells. Upon poly(I:C) challenge, wild-type cells showed a mild increase in PPP1R15A protein expression similar to the *in vivo* sepsis model (**Fig. 1B, Fig. 2C**). In contrast, no-uORF cells had notably high basal expression of PPP1R15A and no further increase was observed after poly(I:C) challenge. Similar patterns were observed in cells harboring easy-to-read uORF (short uORF and no polyproline) (**Fig. 2C, Suppl Fig. 3A-C**). The lack of increases in PPP1R15A protein levels after poly(I:C) in these mutant cell lines was not due to increased protein turnover, reduced PPP1R15A mRNA expression, nor aberrant mRNA splicing (**Fig. 2E, Suppl Fig. 3D, E**). This seemingly paradoxical blunted PPP1R15A expression became even more pronounced when these cells were challenged with higher doses of poly(I:C) (**Fig. 2F, Suppl Fig. 3F**). We reasoned that in these mutant cells, uninhibited PPP1R15A protein expression at baseline could cause excessive translation of antiviral and inflammatory molecules upon poly(I:C) challenge. This in turn can lead to breakthrough phosphorylation of eIF2α despite high level of PPP1R15A, resulting in translation shutdown including the translation of PPP1R15A protein itself. Indeed, we found that cells with easy-to-read uORF (including no uORF) had a prominent antiviral response, underscoring the importance of wild-type uORF in limiting inflammation (**Fig. 2G**). This breakthrough eIF2α phosphorylation in the mutant cells upon poly(I:C) challenge can also explain the relatively small gain in translation (**Fig. 2D and Suppl Fig. 3G bottom**). Surprisingly, long uORF (effectively no CDS expression) also exhibited a heightened antiviral response. This indicates that some degree of PPP1R15A expression is required for preventing excessive phosphorylation of eIF2α and resultant integrated stress response. In summary, PPP1R15A uORF is a major determinant of PPP1R15A protein translation and the removal of the uORF massively increases PPP1R15A protein expression at baseline and under moderate stress. During severe stress, such uninhibited PPP1R15A protein expression does not override eIF2α phosphorylation. Rather, the translation efficiency of PPP1R15A is reduced, underscoring the robust feedback loop system operative along the eIF2α axis^26,27^ (**Fig. 3H**).

### Effects of Ppp1r15a uORF knockout *in vivo*

To determine the significance of Ppp1r15a uORF *in vivo*, we next generated a knock-in mutant mouse model in which the uORF was abolished by introducing a point mutation in its start codon (Ppp1r15a uORF^-/-^, **Fig. 3A**). Mutant mice were born at expected Mendelian ratios with no gross abnormalities (**Suppl Fig. 4A**). Similar to the cell culture model, elimination of Ppp1r15a uORF resulted in a marked increase in basal expression of Ppp1r15a protein in various organs (**Fig. 3B-C**). However, and in contrast to the cell culture model, several organs showed increases in overall translation at baseline, implicating that low grade phosphorylation of eIF2α is operative in these tissues under normal condition (kidney and brain, **Fig. 3D, Suppl Fig. 5A**).

**Figure 4.**
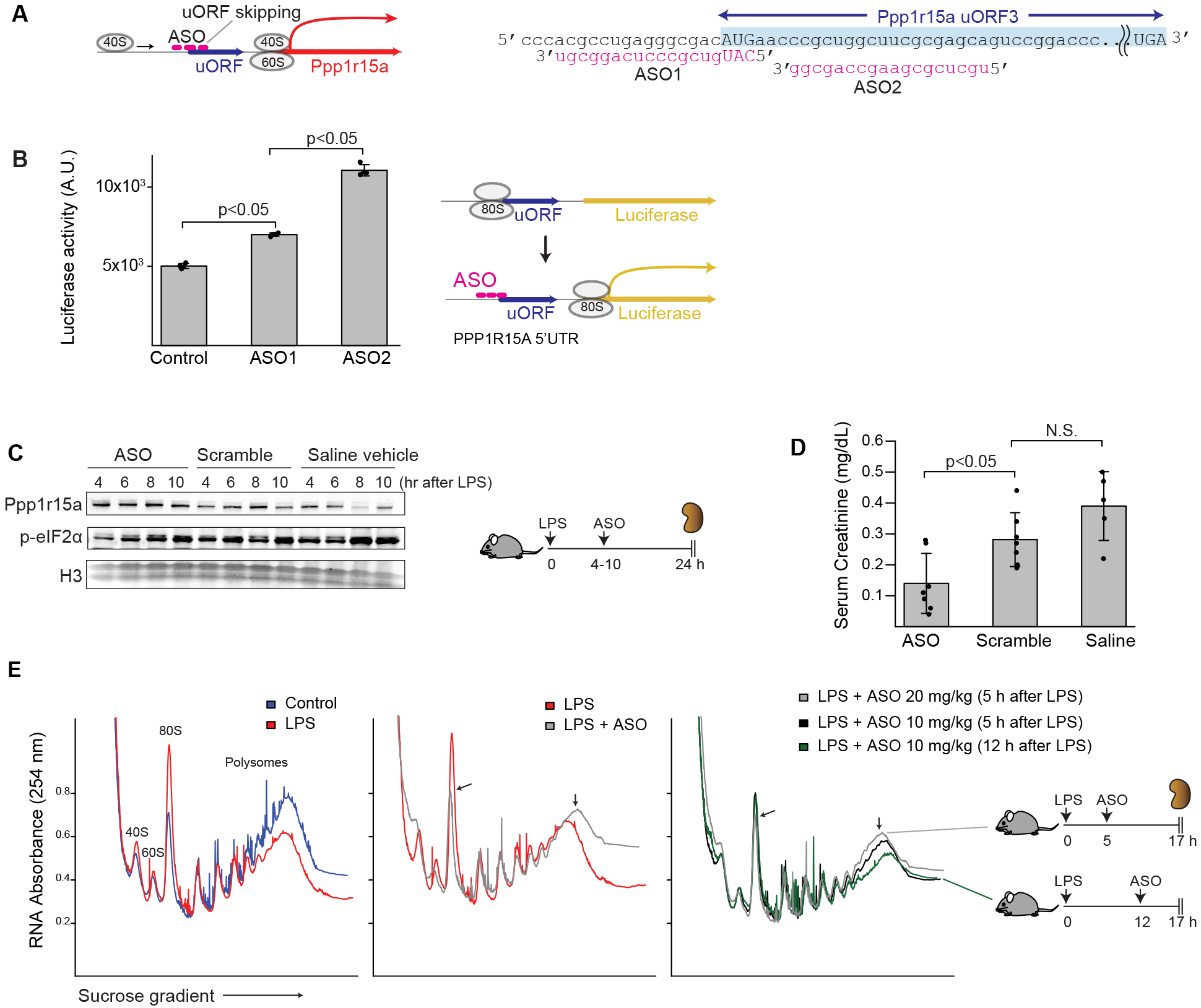
In vivo therapeutic effects of uORF-targeted oligonucleotides in sepsis. (**A**) ASO-based strategy to increase translation efficiency is shown. (**B**) Luciferase activity levels are shown for HEK293T cells transfected with wild-type PPP1R15A 5’UTR luciferase reporter (see **Fig. 1C**) and indicated ASOs. Control consists of scramble ASO mix. (**C**) In vivo time-course experiment demonstrating the upregulation of Ppp1r15a protein levels in the kidneys of ASO-treated mice as determined by Western blot. Data for ASO2 and its scramble are shown. The effect of ASO1 is shown in **Supplemental Figure 7**. Mice were injected with 10 mg/kg ASO via tail vein at indicated time points and tissues were harvested 24 hrs after 5 mg/kg LPS iv. (**D**) Serum creatinine levels 24 hrs after LPS under indicated conditions are shown. ASO2, scramble or saline vehicle were administered one time approximately 8 hrs after LPS. (**E**) Representative polyribosome profiling of kidney extracts from mice treated with indicated conditions. Polysome-to-monosome ratios: 8.4 (control), 4.4 (LPS), 7.4 (LPS + ASO 20 mg/kg 5 h after LPS), 5.2 (LPS + ASO 10 mg/kg 5 h after LPS), and 4.7 (LPS + ASO 10 mg/kg 12 h after LPS).

**Figure 5.**
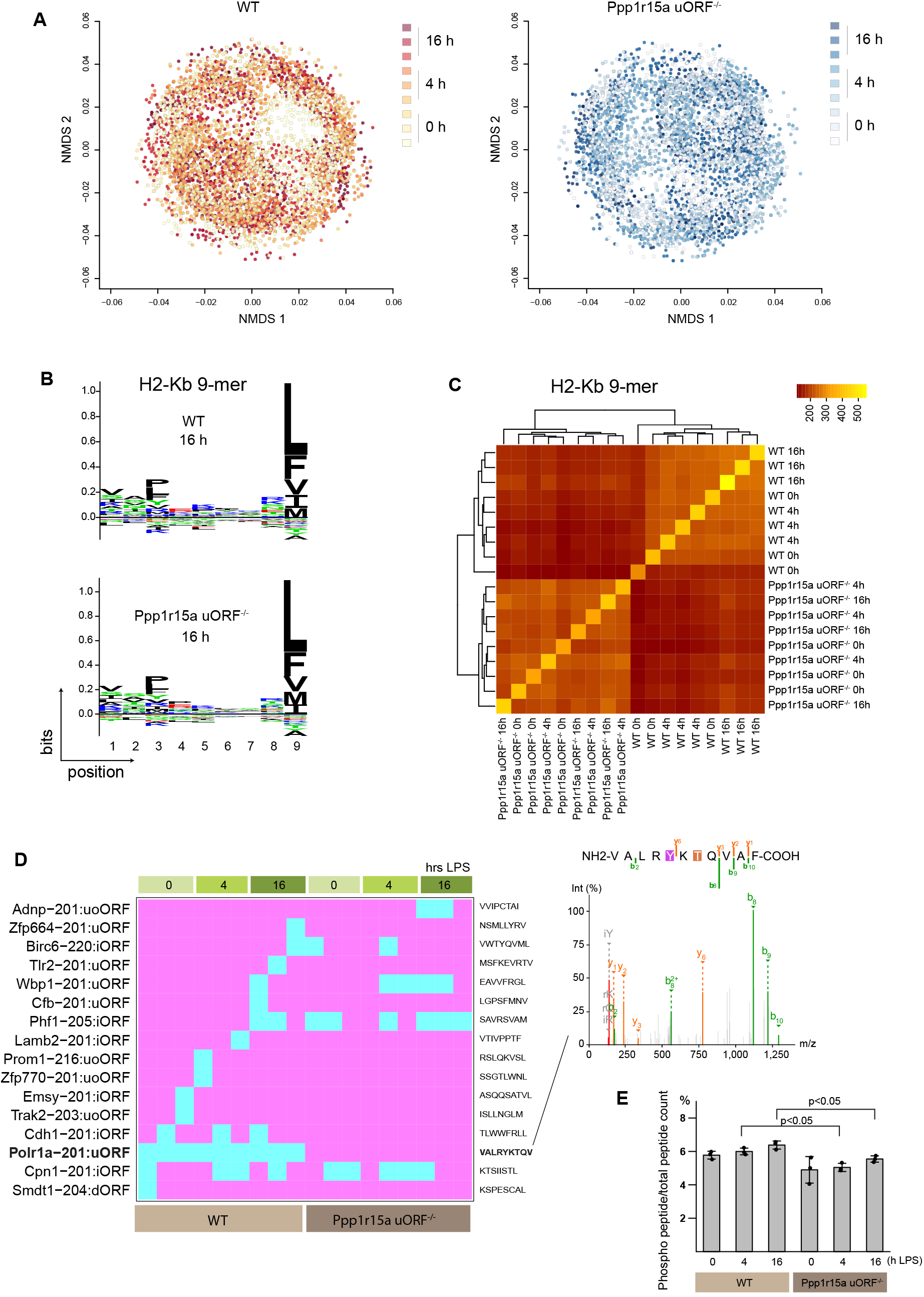
Ppp1r15a uORF^-/-^ mouse kidney exhibits distinct MHC-I immunopeptidome. (**A**) Non-metric multidimensional scaling (NMDS) plots of MHC-I associated peptides detected from WT and Ppp1r15a uORF^-/-^ mouse kidneys. H2-Kb haplotype, 9-mers are shown. Each dot represents a unique peptide. Peptide distance was defined based on amino acid sequence similarity. Overlay of WT and Ppp1r15a uORF^-/-^ NMDS plots are shown in **Supplemental Figure 10**. (**B**) Sequence motif plots are shown for indicated conditions. (**C**) Jaccard similarity index analysis performed on H2-Kb 9-mers per sample. (**D**) Summary of non-canonical peptides supported by both Ribo-seq and MHC-I immunopeptidomics under indicated conditions. LC-MS/MS spectrum of Polr1a uORF is shown on the right. uoORF (upstream overlapping ORF), iORF (internal out-of-frame ORF), dORF (downstream ORF). (**E**) Ratios of phosphorylated peptides adjusted for total peptide counts are shown under indicated conditions.

Time-course analysis of polyribosome profiling after LPS challenge revealed that uORF^-/-^ kidneys did not exhibit the typical early surge in translation that occurs in wild-type. However, and similar to wild-type mice, we noted some degree of translation shutdown in uORF^-/-^ at later timepoints. Nevertheless, overall protein synthesis remained higher in uORF^-/-^ as compared to wild-type, especially at early and late timepoints (**Fig. 3D**). The degree of tissue antiviral responses was comparable between uORF^-/-^ and wild-type kidneys, and renal function was better in uORF^-/-^ animals (**Fig. 3E-F, Suppl Fig. 5 B-D, Suppl Fig. 6**). These findings do not fully agree with the cell culture data, highlighting the complexity of *in vivo* systems. Moreover, we noted that translational changes were tissue dependent. Brain tissue reached its translational nadir earlier than the kidney while overall translation remained elevated in the uORF^-/-^ brain as compared to the wild-type counterpart (**Suppl Fig. 5A**). Unlike the kidney, the liver did show an early surge in translation in uORF^-/-^ but not in wild-type (**Suppl Fig. 5E**). Collectively, these complex features point to the existence of tissue-specific tug-of-war between eIF2α phosphorylation and dephosphorylation as determined by the balance between antiviral response and Ppp1r15a induction, leading to the observed dynamic changes in translation. The finetuning property of the uORF was further confirmed in a model of cecal ligation and puncture in which heterozygous uORF mutation, but not homozygous mutation, led to improved survival, suggesting that mild upregulation of Ppp1r15a offers the optimum balance as a whole in this model of sepsis (**Fig. 3G**).

### Strategy to modulate Ppp1r15a uORF *in vivo*

Having established the significance of Ppp1r15a uORF with genetic approaches, we next developed an *in vivo* uORF knockdown strategy whereby translation rescue could be temporally controlled. To this end, we designed antisense oligonucleotides (ASO) targeting the uORF with the idea that masking of uORF by ASO enables continued ribosome scanning and boosts translation efficiency of downstream CDS (**Fig. 4A**). As inspired by a recent proof-of-concept study^28^, we specifically focused on regions adjacent to the uORF start codon. ASOs were modified with 2’-*O*-methyl to enhance binding affinity and to confer resistance to nucleases. Using luciferase as a readout, we determined that ASO targeting downstream of Ppp1r15a uORF start codon was particularly effective in inducing CDS translation (**Fig. 4B, Suppl Fig. 7A**).

We next tested the efficacy of ASO-based induction of Ppp1r15a protein *in vivo* using our mouse model of endotoxemia. The tissue distribution of ASO is similar to *in vivo* siRNA^29,30^. Indeed, ASO enriches in the proximal tubules^31-33^ where translation shutdown is most prominent in the kidney^12^. We found that the antisense therapy enhanced Ppp1r15a protein expression, improved overall translation, and promoted functional recovery in the kidneys of septic mice (**Fig. 4C-E, Suppl Fig. 7B**). The effects were dose- and time-dependent, and full reversal of eIF2α phosphorylation was not observed with very late intervention (**Suppl Fig. 7C**).

### Effects of Ppp1r15a uORF on MHC-I immunopeptidome

Finally, we sought to identify and characterize a Ppp1r15a uORF-derived peptide. Nearly 50% of our genes harbor at least one uORF and many of them are believed to be translated into peptides based on their ribosome footprints^20,34^. Such peptides could have broad effects beyond modulating scanning ribosome kinetics as determined by uORF sequence characteristics. However, the actual existence of the vast majority of putative uORF-derived peptides await further experimental evidence including a Ppp1r15a uORF-derived peptide. Challenges inherent to detection of uORF-derived peptides include limited tagging, enrichment strategies, and rapid turnover^16,17,35-38^. Indeed, Ppp1r15a uORF contains a degron signal that could cause the peptide to be particularly short-lived (**Suppl Fig. 8A**). We employed a variety of approaches to detect a Ppp1r15a uORF-derived peptide. However, these efforts were met with inconsistent and low confidence results. For example, we failed to detect StrepII-tag-fused uORF peptides by pulldown and mass spectrometry (MS), possibly due to a combination of buried epitope, rapid turnover, and trypsin-digestion of an already short peptide (**Fig. 2A**). We also designed plasmids where additional StrepII-tag (Twin-Strep) was added, and in some experiments, the positions of upstream and downstream Ppp1r15a uORFs were swapped (**Suppl Fig. 8B**). Transfection of these constructs still did not enrich the target peptide with high confidence. Pulldown of UV-crosslinked PPP1R15A mRNA followed by MS revealed multiple RNA-binding proteins (**Suppl Fig. 8C**). However, the target peptide could not be detected. Addition of synthetic uORF-derived peptides into cells or culture medium did not affect its CDS protein levels (**Suppl Fig. 8D-E**). Other strategies we applied are described in Methods (His and FLAG tags fused to N- and C-terminus of the uORF, split fluorescent protein approach, and the use of proteosome inhibitor and endopeptidase inhibitor cocktail). These findings are in line with the literature that detection of uORF peptides is challenging^20,39^.

Notably, Ppp1r15a uORF-derived peptide is predicted to have two protein phosphatase docking motifs (LQPP and FWQP; **Suppl Fig. 8A**). Given that Ppp1r15a is a protein phosphatase regulatory subunit, the enrichment of these protein phosphatase motifs within this 26-mer uORF is highly intriguing. Ab initio protein structure tools^40,41^ predict that Ppp1r15a uORF-derived peptide will form an alpha-helix structure with flexible N-terminus (**Suppl Fig. 8F**). Further computational modeling suggests that this uORF-derived microprotein could interact with protein phosphatase 1 (PP1) near the PP1 catalytic domain^11,42^ (**Suppl Fig. 8G**). We therefore performed PP1 immunoprecipitation (IP) followed by MS. PP1 IP yielded expected enrichment of diverse PP1 binding partners (**Suppl Fig. 8H**). However, we could not detect the target uORF peptide. We further explored potential binding targets of Ppp1r15a uORF peptide using a phage display strategy on synthetic uORF-derived peptides (**Suppl Fig. 9A-B**). Identified potential binding candidates include Lamin B2, Leukocyte receptor cluster member 8, and Inversin/nephrocystin-2 (**Suppl Fig. 9C**).

Recent work by others revealed that uORF-derived peptides are presented on the major histocompatibility complex I (MHC-I)^43,44^. As compared to whole proteome, immunopeptidome (MHC-I IP/MS) significantly enriches non-canonical ORF-derived peptides, possibly due to preferential antigen presentation or stabilization of short non-canonical peptides in the MHC groove (0.1% in whole proteome vs ∼2% in immunopeptidome)^43^. Thus, in our search for Ppp1r15a uORF-derived peptide, we performed immunopeptidomics of the mouse kidney using our endotoxemia model. To enable the detection of unannotated uORF peptides, we built a custom reference database using our kidney Ribo-seq data (see Methods). To enrich C57BL/6 MHC-I haplotype H2-Kb, we used monoclonal antibody produced by Y-3 hybridoma (**Suppl Fig. 10A, Fig 5B**). We show in **Figure 5A** the landscape of kidney immunopeptidomics for wild-type and Ppp1r15a uORF^-/-^ mice at baseline and after endotoxin challenge. We found that the overall difference in immunopeptidomics profile was more pronounced between the two genotypes than with endotoxin-induced changes, pointing to a significant and novel role the uORF plays in immunological governance (**Fig. 5C, Suppl Fig. 10B**). Pathways involved in oxidative phosphorylation and defense against bacterial invasion were enriched in all samples across timepoints (**Suppl Fig. 10C**). Although our target uORF peptide was not detected with this immunopeptidomics approach, various other uORF-derived peptides were identified. Interestingly, a peptide originating from the uORF of Polr1a (RNA polymerase I subunit A) was found in wild-type mouse kidneys, but not in Ppp1r15a uORF^-/-^ (**Fig. 5D**). Ribo-seq data corroborate that Polr1a uORF is translated in wild-type but not in Ppp1r15a uORF^-/-^ (**Suppl Fig. 11A-D**). In addition, MS spectra indicate that this Polr1a uORF peptide is phosphorylated (**Fig. 5D**). While various modifications of MHC peptides have been reported, pathophysiological implications of uORF peptide phosphorylation are largely unknown. It is tempting to speculate that Ppp1r15a uORF^-/-^ (hence increased Ppp1r15a CDS) affected global phosphatase activity, leading to alteration of peptide turnover. Indeed, we found global reduction of phosphorylated immunopeptides across the LPS timepoints in Ppp1r15a uORF^-/-^ mouse kidneys, uncovering the pervasive dephosphorylation property of Ppp1r15a uORF beyond its canonical target eIF2α (**Fig. 5E**).

## Discussion

Here we propose an essential role for the highly conserved Ppp1r15a uORF in the outcomes of sepsis-induced kidney injury. Sepsis is a complex pathological state where multiple pathways are aberrantly activated^45-50^. While these changes are spatially and temporally diverse, one major driver that underlies this dynamic state is the activation of antiviral responses. These in turn lead to the phosphorylation of eIF2α, causing translation shutdown and persistent tissue injury. This converging point offers an attractive route to develop targeted therapy. However, blind manipulation of the core initiation factor eIF2α could lead to unintended consequences such as uncontrolled translation and complete blockade of the integrated stress response. Therefore, we reasoned that it is safer to modulate eIF2α phosphorylation status through Ppp1r15a. One way to alter expression of Ppp1r15a is by targeting its uORF. Indeed, our studies indicate that Ppp1r15a uORF is directly relevant to the underpinnings of translation shutdown, and it is a druggable target using the antisense approach. Our study also reveals that both excessive and insufficient upregulation of Ppp1r15a can result in negative consequences, highlighting the importance of establishing a refined strategy to calibrate this critical node. A grand challenge is then to develop a pragmatic approach to defining the timeline of sepsis and quantitating translation in order to precisely stratify such uORF-based therapy.

Our study revealed the importance of uORF attributes such as length and codon composition in determining CDS translation. Whether these effects are mediated solely by altered ribosome scanning kinetics over the Ppp1r15a mRNA *in cis*, or through much deeper peptide-level network remains undetermined. Small peptides are generally unstable unless dedicated protective mechanisms exist (e.g., insulin). Indeed, very little is known about localization, modification, and lifetimes of uORF-derived peptides. Our target Ppp1r15a uORF encodes multiple prolines. Thus, aside from ubiquitination, other enzymes such as prolyloligopeptidases could potentially be at play^51^, although our attempts to inhibit these enzymes did not lead to improved peptide detection. Regardless, our study reveals strikingly broad biological consequences mediated by this uORF. In particular, the wide-ranging effects on protein phosphorylation suggest that Ppp1r15a uORF modulation has powerful influences even outside of the eIF2α axis. Finally, because thousands of genes harbor ribosome-engaged uORFs that are believed to be functional^20,39,52-57^, our general approach to controlling protein levels through uORF manipulation is highly promising.

### Limitations of the study

This is the first report on the role of Ppp1r15a uORF in sepsis using a newly generated knock-in mouse model and mutant cell lines. This work reveals the intricacy of uORF-dependent Ppp1r15a translation and extends our understanding of Ppp1r15a biology at large^6,23,24,26,58-62^. Future work will involve broader disease models, tissues, and cell types with the goal of fully harnessing the power of uORF modulation. Given the significant changes observed in the MHC-I immunopeptidome, it will be interesting to examine the effects of Ppp1r15a uORF on adaptive immunity using long-term disease models. Whether such uORF-mediated biological effects require the genesis of the purported peptide remains unclear. Addressing these limitations is important for proposing uORF-based therapies to control translation and disease trajectory.

## Methods

### Generation of PPP1R15A uORF mutant cell lines

We designed sgRNA and single-stranded oligo DNA nucleotides (ssODN/HDR, homology directed repair donor oligos), and generated mutant HEK293T cell lines using the CRISPR/Cas9 system (**Table 1**). In all cases, target knock-in and PAM block^63^ (synonymous mutation) were introduced in the vicinity of the double-strand break (±10 bp). Cells were cultured in 10 cm^2^ plates to 70% confluency prior to nucleofection. Approximately 150 × 10^3^ cells (5μL) were mixed with 1.49 μL of ssODN (100 μM) and Cas9 complex consisting of 18 μL SF 4D-nucleofector X solution + supplement1 (Lonza V4XC-2012), 6 μL of sgRNA (30 pmol/μL), and 1 μL of Cas9 2NLS nuclease, S. *pyogenes* (20 pmol/μL, Synthego). Nucleofection was done using Amaxa 4D-Nucleofector X (CM-130 program; Lonza). Cells were seeded in 15 cm plates at various concentrations. Clonal isolation was done manually. Following clonal expansion and genotyping, further subcloning was performed in some cases to ensure purity. On average, the rate of successful biallelic HDR knock-in was approximately 10 – 20%, and the rest of clones showed a mixture of NHEJ and HDR knock-in. Less than 10% of clones showed no evidence of mutation at target sites. DNA extraction was done using Quick DNA Miniprep kit (Zymo Research #D3025). PCR was done using Q5 High-Fidelity DNA polymerase (NEB), PCR primers as listed in **Table 1**, and Monarch PCR Cleanup Kit (NEB, T1030). PCR products were electrophoresed in 2% agarose gel (TopVision Agarose Tablets ThermoFisher R2801) and bands were excised and extracted using QIAQuick Gel Extraction kit (Qiagen, #28706). Sanger sequencing was done at GeneWiz. Software used for design and analysis of mutant cell lines include CRISPRdirect, SnapGene, Primer3Plus, NEB Tm calculator, and Synthego ICE.

**Table 1.**
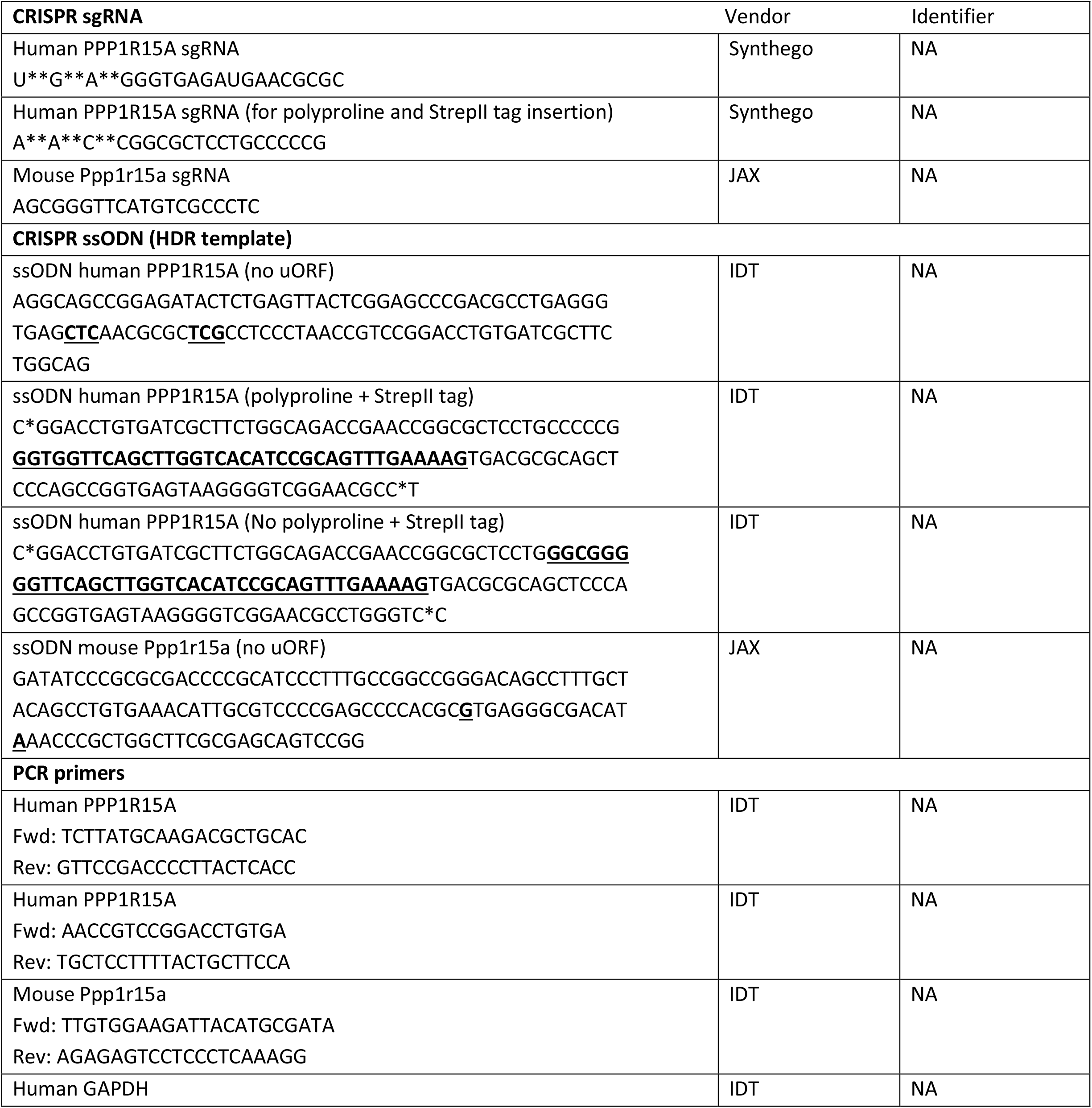
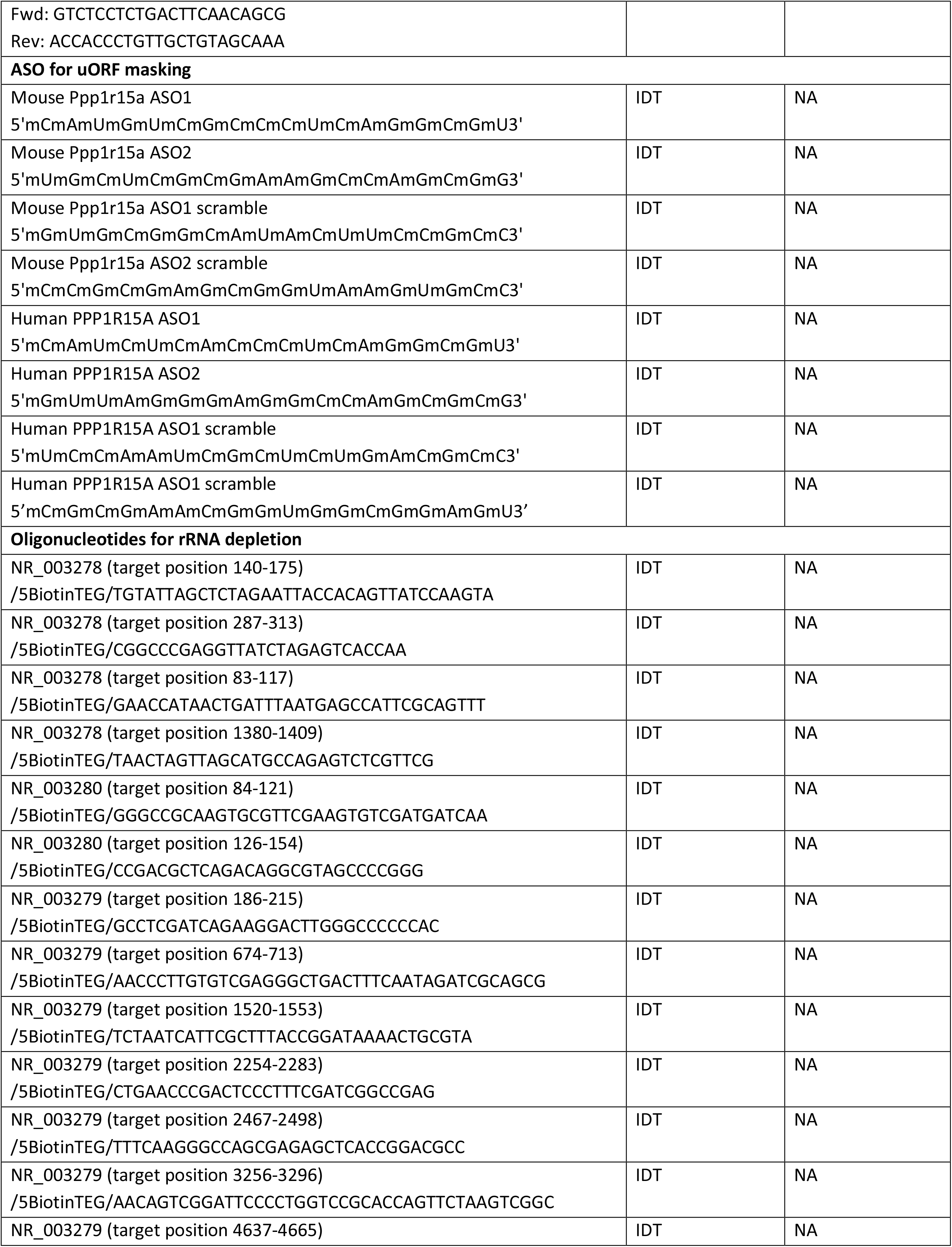
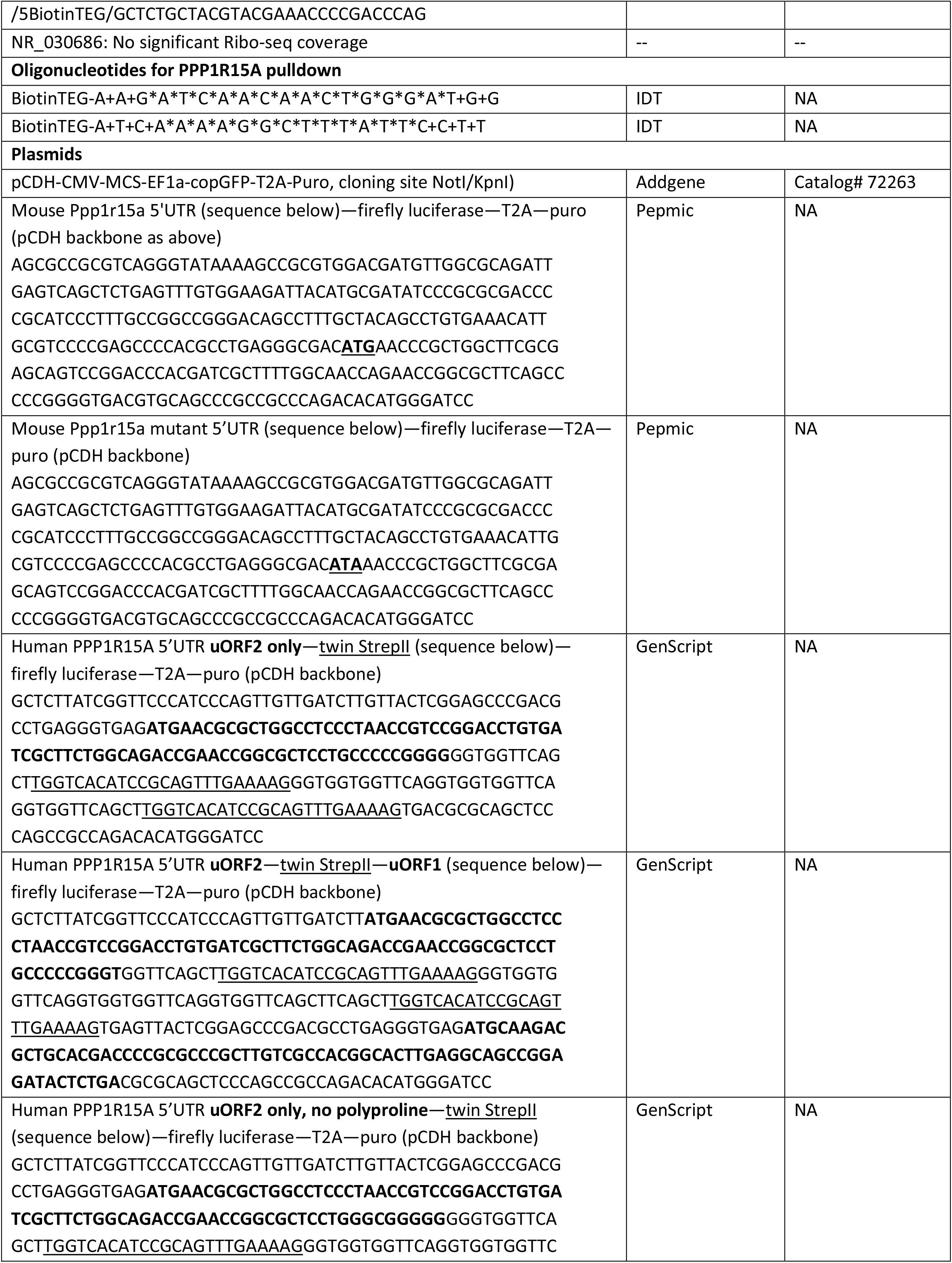
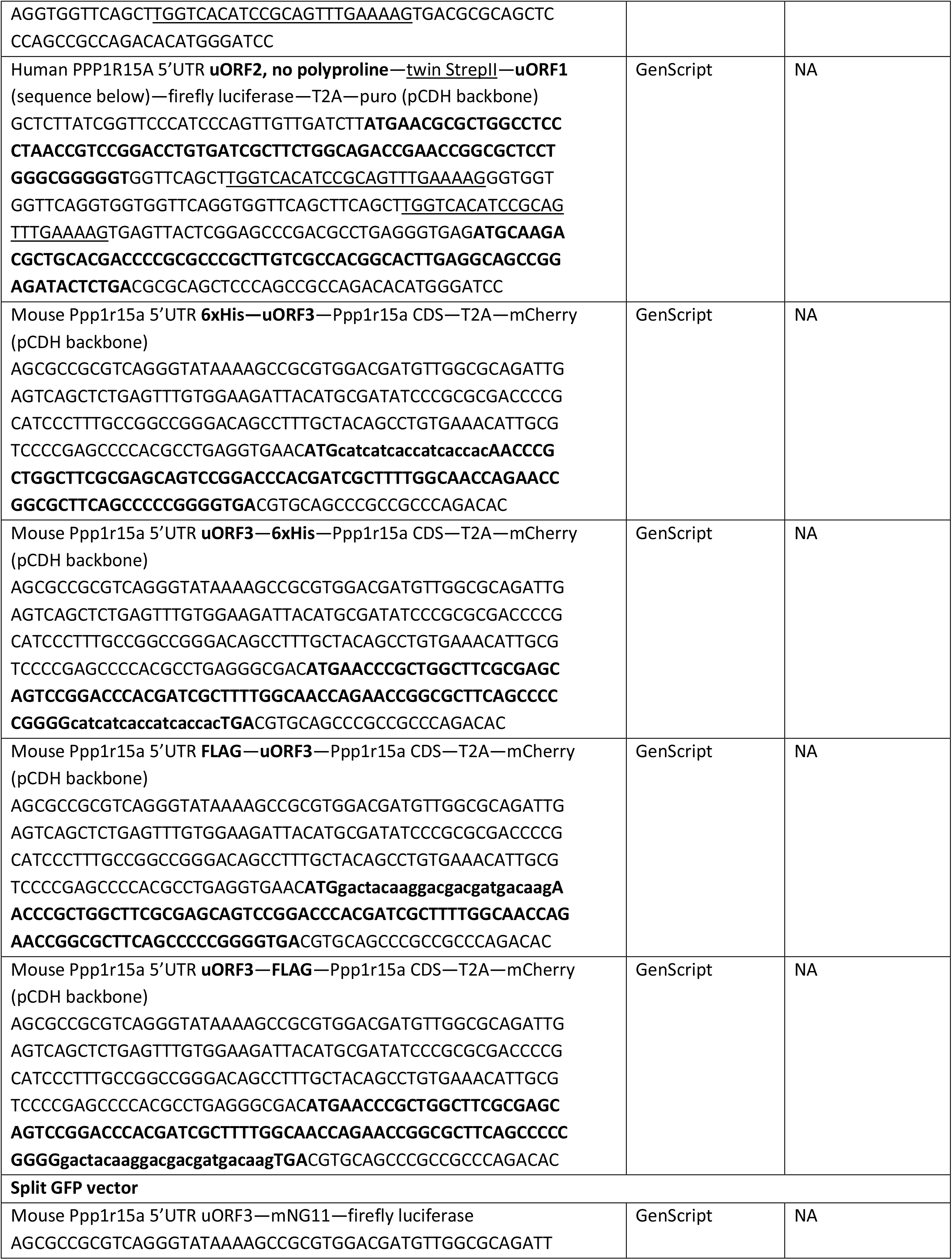
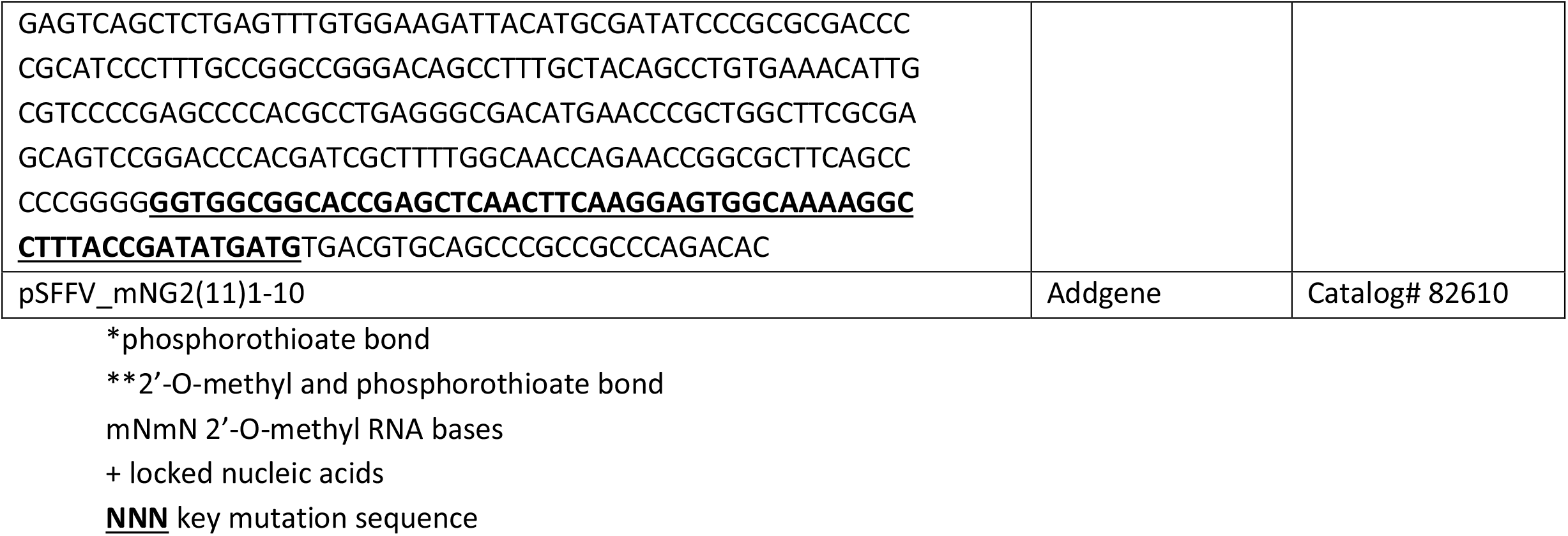
Oligonucleotides.

### Generation of Ppp1r15a uORF mutant mouse model

We designed the following sgRNA and ssODN to introduce uORF3 start codon mutation and PAM block^63^ within 10 bp from the double-strand break. Asymmetric donor DNA strategy^64^ was used (36bp|cut|91bp for the nontarget strand). sgRNA: 5’AGCGGGTTCATGTCGCCCTC3’(AGG/PAM) ssODN/HDR donor oligos: 5’GATATCCCGCGCGACCCCGCATCCCTTTGCCGGCCGGGACAGCCTTTGCTACAGCCTGTGAAACATTGCGTCCCCGAGCCCCACG**<u>C GT</u>**GAGGGCGAC**<u>ATA</u>**AACCCGCTGGCTTCGCGAGCAGTCCGG3’

Embryo manipulation and generation of founder mice on C57BL/6J background were performed by the Jackson Laboratory Mouse Model Generation Services. N1 sperms (heterozygous) have been cryopreserved at the Jackson Laboratory (identifier: IU_GET5678).

### Animals

Ppp1r15a uORF knock-in mice (uORF^-/-^, uORF^+/-^, and uORF^WT^ littermates) were housed at Indiana University School of Medicine under a 12 h light/dark cycle at 25 °C. For studies that did not require the knock-in mice, C57BL/6J mice were obtained from the Jackson Laboratory. All mice were 8–12 weeks of age and weighed 24–32 g. Male mice were used unless indicated. Animals were subjected to a single-dose, 5 mg/kg LPS tail vein i.v. injection (*E. coli* serotype 0111:B4, MilliporeSigma). Untreated mice received an equivalent volume of sterile, normal saline vehicle. Cecal ligation and puncture was performed under isoflurane anesthesia; 75% of the mouse cecum was ligated and punctured twice with a 25-gauge needle. Animals were resuscitated with 500 μl of normal saline i.p. at the time of abdominal closure. No antibiotics were used. For *in vivo* ASO treatment, Ppp1r15a uORF-targeted oligonucleotides or corresponding scrambles (**Table 1**) were reconstituted in water (1 mM stock solution) and rediluted in sterile normal saline with final volume of 300 μL for tail vein i.v. injection.

### Cells

HEK293T cells and PPP1R15A uORF mutant cells were cultured in Dulbecco’s Modified Eagle Medium with 10% fetal bovine serum and 100 U/mL penicillin and 100 μg/mL streptomycin (ThermoFisher). All cell types were cultured at 37 °C with 5% CO_2_.

### Transformation of E. coli and transfection of HEK293T cells

Chemically competent E. coli (One Shot Top10, Invitrogen C404003) were transformed with pCDH plasmids (**Table 1**) using the heat shock method (on ice for 30 min followed by 42 °C heat shock for 45 seconds). Transformed E. coli were incubated overnight in LB broth with 100 μg/mL carbenicillin at 225 rpm, 37 °C. Plasmid isolation was done using Nucleobond Xtra Midi Kit (Takara #740410.50) following the manufacture’s protocol. For DNA gel electrophoresis, 1 % agarose gel was prepared using TopVision Agarose Tablets (ThermoFisher R2801) and Gel green nucleic acid stain (Biotium #41005). Samples were electrophoresed with NEB X6 gel loading dye in 0.5X TAE buffer at 100 V. Transfection of plasmids was done using lipofectamine 2000 following the manufacturer’s instruction (sequential mixing of lipofectamine, Opti-MEM and plasmid into freshly replaced DMEM/10% FBS medium, ThermoFisher 11668027). For all experiments, either 1 μg or 10 μg plasmid DNA was used per well of a 6 well-plate or 10 cm dish, respectively. For split GFP co-transfection, 1 μg of mNG2(11)1-10 and 1 μg of Ppp1r15a 5’UTR uORF3—mNG11 were added per well of a 6 well-plate. GFP signal could not be detected with overnight co-transfection (data not shown).

Poly(I:C) transfection: Cells with 60-80% confluency were transfected with synthetic double-stranded RNA poly(I:C) (high molecular weight, InvivoGen #tlrl-pic) using lipofectamine 2000 following the manufacture’s protocol. Final working concentrations of poly(I:C) ranged from 0.1 μg/mL to 1.5 μg/mL. Cells were incubated for 1 to 16 hours and harvested for downstream analyses. In some experiments, other stress-inducing reagents were used (without transfection): tunicamycin 10 μg/mL (Sigma), thapsigargin 1 μM (Sigma), recombinant human interferon gamma protein (20 ng/mL, R7D Systems 285-IF).

For ASO transfection, Lipofectamine RNAiMAX (ThermoFisher) was used following the manufacture’s protocol (80 nM of indicated ASO for 6 hrs). Transfection of synthesized uORF peptide was done using Chariot Protein Delivery Reagent (ActiveMotif #30025) following the manufacture’s protocol. Fifteen μM of uORF peptide complexed with Chariot reagent was transfected and incubated for 2 hrs before cell lysis. In a separate experiment, 1.5 μM of the peptide was added to freshly replaced culture medium and incubated for 2 hrs.

### Luciferase assay

HEK293T cells were split on previous day to result in a 60-80% confluency. Cells were transfected with 10 μg of pCDH-Ppp1r15a 5’UTR (uORF3 start codon mutant)-luciferase-T2A-puroR or pCDH-Ppp1r15a 5’UTR (WT)-luciferase-T2A-puroR constructs per 10 cm petri dish with lipofectamine 2000 overnight. Cells were split into 96-well plates at 50,000 cells/well in 200 μL DMEM without penicillin/streptomycin and allowed to adhere overnight. Detection of luciferase signal was done using the Bright Glo luciferase assay system (Promega, E2610). Following cell incubation at 37°C, luciferase luminescence signal was read in Clario Start plate reader.

### Polyribosomal profiling

For polyribosome profiling of tissues, cardiac perfusion was performed with 6 mL of cycloheximide (100 μg/ml in PBS, Sigma). Harvested tissues were immediately placed in a lysis buffer consisting of 1% Triton X-100, 0.1% deoxycholate, 20 mM Tris-HCl, 100 mM NaCl, 10 mM MgCl_2_, EDTA-free Protease Inhibitor Cocktail Tablet (Roche) and 100 μg/ml cycloheximide. Tissues were homogenized using a Minilys tissue homogenizer (Bertin Instruments). Tissue homogenates were incubated on ice for 20 minutes, then centrifuged at 9,600 *g* for 10 minutes. The supernatant was added to the top of a sucrose gradient generated by BioComp Gradient Master (10% sucrose on top of 50% sucrose in 20 mM Tris-HCl, 100 mM NaCl, 5 mM MgCl_2_, and 100 μg/ml cycloheximide) and centrifuged at 283,800 *g* for 2 hours at 4 °C. The gradients were harvested from the top in a Biocomp harvester (Biocomp Instruments), and the RNA content of eluted ribosomal fractions was continuously monitored with UV absorbance at 254 nm. For polyribosome profiling of cultured cells, cells were rinsed with cold PBS first, then cells were scraped using the lysis buffer on ice. The remaining procedure was identical to tissue polyribosome profiling.

### Ribo-Seq analysis

Since we reported our prior ribo-seq analysis^12^, Illumina TruSeq RNase was discontinued. Consequently, we reoptimized our workflow primarily focusing on the choice and condition of RNase digestion. RNase T1, S7 micrococcal nuclease, and RNase I were evaluated using polyribosomal profiling (enrichment of monosomal peak and disappearance of polysomes as a readout)^65^ and sequencing. We opted for RNase I because it has no sequence bias in its endonuclease activity. However, as reported by others^18^, we also noted that ribosome footprint periodicity was not fully preserved with RNase I, possibly due to the aggressive nature of the enzyme. We also compared custom oligo-based rRNA depletion versus no rRNA depletion since Ribo-Zero Gold (Illumina) was discontinued as a standalone product^66^ (**Table 1**; target sites were determined based on preliminary Ribo-seq without rRNA depletion). While rRNA was depleted with the custom biotinylated oligos, the overall loss of ribosome protected fragments was not negligible. Therefore, we did not include the depletion step in the following workflow.

Kidneys were snap-frozen and pulverized under liquid nitrogen. Cryo-lysis was done using a Minilys tissue homogenizer in a polysome buffer (1 mL per one mouse kidney) consisting of 20 mM Tris pH 7.5 (140 μL of 1 M Tris 7.0 plus 60 μL of 1 M Tris 8.0 for 10 mL final buffer), 100 mM NaCl, 10 mM MgCl_2_, 1% Triton X-100, 0.1% sodium deoxycholate (Sigma D-6750), DTT, 0.02 U/μL Turbo DNAse (ThermoFisher AM2238), 200 μg/mL cycloheximide (Sigma), and Protease Inhibitor Cocktail Tablet (Roche). Tissue lysates were centrifuged at 20,000 *g* for 10 minutes. Equal volumes of supernatant aliquots were made for ribo-Seq and RNA-Seq workflows (200 μL each). Polysome digestion was performed using RNase I (ThermoFisher/Ambion AM2294) for 60 minutes at 4 °C and the reaction was stopped with 10 μl SuperaseIn (Thermo Fisher Scientific AM2696). Ribosome-protected fragments were isolated using microspin S-400 (GE HealthCare) following the manufacturer’s instruction. Microspin buffer consisted of Tris pH 7.4 20 mM, NaCl 150 mM, MgCl_2_ 5 mM. RNA fragments were purified using the TRIzol/chloroform method and chill precipitated in 3M sodium acetate and GlycoBlue overnight at -20 °C. We next performed time-gated size selection using the Pippin HT system (7:00 min to 28 min on 3% agarose; Sage Science). Size selected RNA samples were purified using Zymo RNA Clean & Concentrator-5 kit (Zymo R1013). RNA 3’ end dephosphorylation reaction consisted of T4 PNK (20 U/10 μL sample, NEB M0201S), SuperaseIn in 1X T4 PNK buffer without ATP for 60 min at 37 °C. Adapter ligation was done using NEXTFLEX Small RNA-seq kit v3 (PerkinElmer NOVA-5132-05). After 3’ 4N adenylated adapter ligation and excess adapter inactivation (STEP C in the NEXTFLEX manual), 5’ end repair was done using T4 PNK with 2 mM ATP (NEB P0756S). Following RNA purification with Zymo Oligo Clean & Concentrator (D4060), we proceeded with the remaining NEXTFLEX steps through G following the manufacture’s manual. Library size selection for sequencing was done using the Pippin System (Range mode, 135 bp to 190 bp). Sequencing was performed at Indiana University School of Medicine Medical Genomics Core using Illumina NovaSeq (single-end 75 bp reads). Cutadapt was used to trim adapters including the 4 random oligonucleotides between each adapter and sample read. Reads were first aligned to rRNA followed by mapping to the rest of the mm10 reference genome using STAR and in some cases bowtie for transcriptome. Data analysis was done using multiple tools we previously described^12^. Gedi PRICE^67^ was used for detection of ATG and non-ATG uORF genome-wide, and construction of a custom reference for MHC peptidomics. For determination of reads coverage, P-site, and periodicity, samtools, bedtools, awk, and the following R packages were used: RiboWaltz, GenomicRanges, IRanges and BiocGenerics.

### RNA-Seq

Kidneys were snap-frozen and RNA was extracted using Zymo Direct-zol RNA kit. RNA quality was determined using Agilent Bioanalyzer (RIN values > 7). Sequencing was done at the Indiana University Center for Medical Genomics Core. One hundred ng of total RNA was used. cDNA library preparation included mRNA capture/enrichment, RNA fragmentation, cDNA synthesis, ligation of index adaptors, and amplification, following the KAPA mRNA Hyper Prep Kit Technical Data Sheet, KR1352 – v4.17 (Roche). Each resulting indexed library was quantified, and its quality accessed by Qubit and Agilent Bioanalyzer, and multiple libraries pooled in equal molarity. The pooled libraries were then denatured, and neutralized, before loading to NovaSeq 6000 sequencer at 300 pM final concentration for 100 bp paired-end sequencing (Illumina). Approximately 40M reads per library was generated. Phred quality score (Q score) was used to measure the quality of sequencing. More than 90% of the sequencing reads reached Q30 (99.9% base call accuracy). The sequenced data were mapped to the mm10 genome using STAR. Uniquely mapped sequencing reads were assigned to mm10 refGene genes using featureCounts and counts data were analyzed using edgeR.

### MHC peptidomics

To harvest a large amount of monoclonal antibody against MHC-I haplotype H2-Kb, Y-3 hybridoma cells (ATCC HB-176) were incubated in a membrane cell culture flask following the manufacturer’s instructions (Wheaton CELLine Bioreactor Flask and Hybridoma-SFM ThermoFisher 12045076). Crosslinking of Y-3 antibody and Protein A+G was done using sulfosuccinimidyl (4-iodoacetyl)aminobenzoate (Sulfo-SIAB, ThermoFisher 22327). For 1 mL of protein A+G agarose (ThermoFisher/Pierce #20421), 1.5 mL of Y-3 antibody (0.5 mg/mL) was added and incubated at room temperature for 40 min. Next, Sulfo-SIAB (25 mM working concentration) was added and incubated for 30 min in dark. Excess crosslinkers were removed by washing with 50 mM borate buffer (50 mM borate buffer, 5 mM EDTA, pH 8.5). Crosslinking reaction was quenched with 5 mM cysteine incubation at room temperature for 15 min in dark. Crosslinked column was then washed with PBS. Mouse kidney tissues were lysed using Minilys (45 seconds at the highest speed) with the following buffer: 20 mM Tris pH 8, 1 mM EDTA, 100 mM NaCl, 1% Triton X-100, 60 mM Octyl Glucoside (Sigma), 2 mM MgCl_2_, DNase I (0.1 U/μL, Ambion AM2222), Benzonase (25 U/mL EMDMillipore 70746-4), 10 mM chloroacetamide (Sigma), cOmplete Protease Inhibitor Cocktail (Sigma), and phosStop inhibitor (Roche). Samples were left on ice for 10 min and centrifuged at 12,000 rpm for 10 min. The supernatant was mixed with crosslinked Y-3 antibody and incubated for 2 hrs at 4 °C. The resin was washed with 10 column volume wash buffer (10 mM Tris pH 8.0, 1 mM EDTA, 100 mM NaCl, 2 mM MgCl_2_, and 0.1% Triton X-100), 10 column volume 10 mM Tris (pH 8), followed by water. MHC peptides were eluted in 0.1 N acetic acid for 2 hrs, and desalted on 50 mg SepPak columns (Waters) with 0.5% trifluoroacetic acid. After washing with 0.1% trifluoroacetic acid, MHC peptides were eluted in 30% and 60% acetonitrile with 0.1% formic acid. Lyophilized samples were reconstituted in 0.1% formic acid prior to LC-MS/MS analysis on Orbitrap Eclipse mass-spectrometer with FAIMS Pro Interface (Thermo Scientific). The LC linear gradient consisted of 6-30% Buffer B (0.1% formic acid and 90% acetonitrile) over 84 min, 30-90% Buffer B over 9 min and held at 90% Buffer B for 5 min at 200 nl/min following the protocol by Ouspenskaia et al^43^. (Buffer A: 0.1% formic acid). Data were analyzed in MSFragger/FragPipe using the default option for MHC peptidomics (“Nonspecific-HLA”, closed search)^68^ except for peptide lengths, which were restricted to 8 – 12 amino acids, and addition of STY phosphorylation as a variable modification. Reference FASTA file was prepared by combining the UniProt mouse reference (UP000000589_10090), common contaminants, custom uORF database (see below), and decoys (reverse sequence). Only top hit-rank peptides (hit rank=1) were subsequently analyzed using NetMHCpan4.1b for H2-Kb and H2-Db with default settings (Rank threshold for strong binding peptides 0.5, weak binding peptides 2.0)^69^. Peptides with binding %rank < 2.0 were further filtered for 9-mers and H2-Kb. Non-metric multidimensional scaling was done using R packages motifStack, HDMD and ecodist as described by Sarkizova et al^70^. Nine-mer logo display was done using seq2logo. Jaccard analysis was done using base R function %in% and heatmap3. Pathway analysis was done using pathfindR.

Regarding custom uORF database, we built it based on kidney Ribo-seq data^12^ in which a model of endotoxin-induced kidney injury was applied similar to the peptidomics study. This allowed us to construct a condition-specific uORF database. Because non-ATG start codon use is pervasive in uORF, computational ORF prediction (canonical ATG start|TGA/TAA/TAG stop) does not capture a full spectrum of translated uORF. Thus, to systematically detect uORFs from Ribo-seq data, we used Gedi PRICE pipeline^67^. BAM files were converted to CIT files and detected ORFs were annotated according to PRICE ORF annotation types (https://github.com/erhard-lab/gedi/wiki/Price). uoORF, uORF, iORF, and dORF reads from the main output table (orfs.tsv) were reorganized and converted to a BED file. Transcription and translation of the BED file were done using bedtools (getfasta function) and faTrans.

### Immunoprecipitation and mass spectrometry

Cells were lysed using Minilys homogenizer with the following extraction buffer: 20 mM Tris pH 8, 100 mM NaCl, 1% Triton X-100, 2 mM MgCl_2_, DNase I (0.1 U/μL, Ambion AM2222), Benzonase (25 U/mL EMDMillipore 70746-4), Halt protease inhibitors (Pierce), and phosStop inhibitor (Roche). For PP1 IP, supernatants were incubated with anti-PP1 antibody conjugated to agarose for 3 hrs at 4 °C (E-9, sc-7482 AC, 25% agarose, Santa Cruz). Samples were washed with washing buffer 5 times (20 mM Tris pH 8, 500 mM NaCl, 0.1% TritonX 100 and 2 mM MgCl_2_). Samples were trypsin digested, desalted, and run on Orbitrap Exploris 480 mass spectrometer with FAIMS. Pulldown of StrepII tag was done following the MagStrep-Tactin beads protocol (IBA-Lifesciences, 2-4090-002). For pulldown of PPP1R15A mRNA and RNA-bound proteins, we followed the protocol described by Iadevaia et al^71^. Cells were UV-crosslinked and lysed in a buffer consisting of 100 mM Tris-HCl, pH 7.5, 500 mM LiCl, 10 mM EDTA, 1% Triton X-100, 5 mM DTT, 20 U/ml DNase I, SuperaseIn, cOmplete Protease Inhibitor Cocktail. 5’ biotin-TEG modified antisense oligos targeting the 5’ and 3’ ends of PPP1R15A mRNA were prepared (**Table 1**) and incubated at 70 °C for 5 min. Supernatants were incubated with the denatured oligos for 30 min at 25 °C, then streptavidin-coupled magnetic beads were added (Dynabeads MyOne Streptavidin C1, ThermoFisher 65001). RNA and proteins were recovered following the Iadevaia’s protocol^71^. In a separate experiment (without immunoprecipitation), cells were treated with poly(I:C) (1.5 μg/mL for 4 hrs) and a combination of proteasome inhibitor (Bortezomib, 2 μM, Sigma/Calbiochem 5043140001) and prolyl endopeptidase inhibitors (Pramiracetam 10 uM, Sigma SLM-0816; Baicalein, 10 uM, Sigma 465119; Berberine Chloride Sigma 10 uM, B3251) for 4 hrs, and cell lysates were analyzed using Orbitrap Exploris 480 mass spectrometer.

### Western blotting

Proteins from cells and tissues were extracted using RIPA buffer (ThermoFisher Pierce) with 0.5M EGTA, 0.5M EDTA, DNase I (0.1 U/μL, Ambion AM2222), Halt protease inhibitors (Pierce), phosStop inhibitor (Roche), and benzonase nuclease (25 U/mL EMDMillipore 70746-4). Total protein levels were determined using a modified Lowry assay (Bio-Rad). Equal amounts of total proteins (10 - 20 μg) were mixed with NuPAGE LDS Sample Buffer (Thermo Fisher) with 100 mM of DTT and separated by electrophoreses on NuPage 4%–12% Bis-Tris gels and transferred to PVDF membranes. Antibodies used include the following: Ppp1r15a/GADD34 (Proteintech, rabbit polyclonal, 1:1,000 dilution, #10449-1-AP), p-eIF2α (Ser51, Invitrogen, catalog PIPA537800), eIF2α (Cell Signaling Technology, catalog 9722), mIgG2b kappa isotype (control antibody for Y3 antibody western blot, 1:3,000 dilution, eBioscience/ThermoFisher #14-4732-82), Alexa Fluor 680 and 790 secondary antibodies (Santa Cruz Biotechnology), and histone H3 (Cell Signaling Technology, mouse monoclonal, 1:1,000 dilution, catalog 9715). For PPP1R15A protein turnover determination, cells were incubated with 250 μg/mL of cycloheximide for indicated duration prior to cell lysis.

### PCR

RNA was extracted using TRI Reagent and Direct-zol RNA MiniPrep Plus (Zymo Research R2070). Reverse transcription was done using High Capacity cDNA Reverse Transcription kit (ThermoFisher/Applied Biosystems 4368814). For SYBR Green qPCR, SensiFAST SYBR Green No-ROXSYBRgreen (Bioline BIO-94002) was used on ViiA7 Real-Time PCR systems (ABI). The ΔΔCt method was used to analyze the relative changes in gene expression. PCR primers used are listed in **Table 1**. Conventional PCR was done using 2% agarose gel.

### Peptide synthesis and phage display

The uORF-derived peptide synthesis and phage display library screening were done at the Chemical Genomics Core facility in Department of Biochemistry and Molecular Biology, Indiana University School of Medicine. The peptide was synthesized by standard solid phase peptide synthesis with biotin and spacer conjugated to the N terminal (**Supplemental Figure 9**). High-resolution mass spectra and purity data were obtained using an Agilent 1290 LC -6545 QTOF. The purity of the peptide was established to be over 90% (UV, λ = 254 nm) and HRMS (ESI): (M+2H)^+2^ m/z calculated for: 1723.9934, found: 1723.8217. The biotinylated peptides were placed on streptavidin-coated plates and three rounds of solid-phase panning were conducted using 10^9^ random 12-mer peptides fused to a minor coat protein of M13 (pIII) following the manufacture’s protocol (Ph.D-12 Phage Display Peptide Library Kit, NEB, E8110S). Sequenced data were subsequently analyzed as follows. Homology modeling of uROF-derived peptide was done using I-TASSER homology modeling server. Modeling of peptide sequences identified was done using de novo structure prediction server PEP-FOLD3. Docking was done using blind peptide-protein docking server HPEPDOCK. Docked models with lowest energy were further passed on to protein binding energy prediction server PRODIGY on the basis of Gibb’s free energy and dissociation constant at 25°C, and visualization was done using PyMOL2.0.

### Immunohistochemistry

Tissues were fixed with 4% paraformaldehyde, sectioned (5 μm), and deparaffinized. Low pH antigen retrieval was used. Tissues were stained for Ppp1r15a (Proteintech, #10449-1-AP) and imaged using a Keyence BZ-X810 fluorescence microscope.

### Single-cell ATAC-sequencing

Single nuclei isolation was done following the Humphreys lab protocol^72^. Each mouse kidney was minced and homogenized using Dounce homogenizer (5x loose, 10x tight, Kimble Chase KT885300-0002) in 2 mL of Nuclei EZ lysis buffer (Sigma N-3408) supplemented with SuperaseIn and cOmplete ULTRA Tablets Mini (Roche, 05892791001). Samples were rediluted in the lysis buffer, filtered through a 40-μm cell strainer, and centrifuged for 5 min at 500 g. The pellet was resuspended in 3.5 mL of lysis buffer. After another centrifugation, the pellet was resuspended in Nuclei Buffer provided by 10X Genomics, and nuclei counts were adjusted with additional filtering and centrifugation. The single cell ATAC analysis was conducted using a 10X Chromium single cell system (10X Genomics) and a NovaSeq 6000 sequencer (Illumina) at the Indiana University School of Medicine Center for Medical Genomics Core. Final nuclei concentration of 3000/μL or higher was applied to targeting nuclei recovery of 10,000. Tagmentation was performed immediately, followed by GEM generation and barcoding, and library preparation using Chromium Single Cell ATAC Reagent Kits User Guide, CG000168 Rev A (10X Genomics). The quality of the library was examined by Bioanalyzer and Qubit. The resulting library was sequenced for 50 bp paired-end sequencing on Illumina NovaSeq 6000. About 50K reads per nuclei were generated and 91% of the sequencing reads reached Q30 (99.9% base call accuracy). Phred quality score (Q score) was used to measure the quality of sequencing. Data processing and visualization were done using the Cell Ranger ATAC pipeline and Loupe Brower.

## Data availability

RNA-seq and Ribo-seq data will be deposited in the NCBI’s Gene Expression Omnibus database (GEO GSExxxx). Proteomics data will be deposited in ProteomeXchange (xxxx).

## Code availability

Scripts will be made available through GitHub: https://github.com/hato-lab

## Study approval

All animal protocols were approved by the Indiana University Institutional Animal Care Committee and conform to the NIH *Guide for the Care and Use of Laboratory Animals*, National Academies Press.

## Acknowledgements

We thank Amber Mosley, Aruna Wijeratne and Guihong Qi at the at the Proteomics Core, and Yunlong Liu, Hongyu Gao, Patrick McGuire and Mandy Bittner at the Center for Medical Genomics at the Indiana University School of Medicine, and former lab member Thomas W. McCarthy. We also thank Constance Temm and Connor Gulbronson for assistance with tissue staining and imaging. Measurement of serum creatinine concentration was performed by John Moore, Yang Yan, et al. at the University of Alabama at Birmingham/UCSD O’Brien Center Core for Acute Kidney Injury Research (NIH P30DK079337) using isotope dilution LC-MS. This work was supported by NIH grants R01-AI148282, K08-DK113223, Veterans Affairs Merit (BX002901), Indiana Clinical and Translational Sciences Institute to TH, and R01-DK107623 to PCD.

## Figure legends

**Supplemental Figure 1**

(A) Schematic of the eIF2 axis. Only pertinent kinase (Eif2ak2/PKR) is shown for clarity. (**B**) Mouse kidney Ribo-seq data analysis. P-site offset was computed from ribosome-protected mRNA fragments mapped to transcriptome genome-wide (inset). P-site reads coverage is shown for Ppp1r15a and Ppp1r15b, and these reads are color coded based on their codon frame use. The codon-frame periodicity was calculated from the transcription start site (TSS) in the genome coordinate. Thus, codon frame colors can be different before and after an intronic region for a given coding sequence (CDS). Regular RNA-seq reads are color coded in gray. Note the low translation efficiency of Ppp1r15a (but not Ppp1r15b) as determined by the low ribo-seq-to-RNA-seq ratio over the CDS. Arrowhead in the left upper panel points to distinct 3-nulceotide periodicity throughout the 26 amino acid codons of the 3^rd^ Ppp1r15a uORF (blue). (reanalysis of published data^12^) (**C**) Single-cell RNA-sequencing of mouse kidneys showing transcriptional increases of Ppp1r15a in a wide range of kidney cell types after LPS challenge (reanalysis of published data^13^). (**D**) Single-nuclear ATAC-sequencing (Assay for Transposase-Accessible Chromatin) of mouse kidneys demonstrating the accessibility of Ppp1r15a promoter region in all cell types at baseline and after LPS challenge (red vertical lines). Inset is a summary of transcription factors identified by ChIP-seq at the Ppp1r15a promoter region (ChIP-Atlas, filtered for mouse kidneys and the significance threshold is set at 200). (**E**) Human, mouse and rat Ppp1r15a transcripts are aligned and annotated.

**Supplemental Figure 2**

Sanger sequencing chromatograms for all cell lines used in this study. Key features and mutations introduced by CRISPR/Cas9 are annotated.

**Supplemental Figure 3**

(**A**) Quantitation of PPP1R15A mRNA levels as determined by real-time qPCR under indicated conditions. PCR primers used are identical to the one shown in main **Figure 2C**. n=3. (**B**) Quantitation of PPP1R15A protein levels as determined by western blot under indicated conditions. n=3, independent replicates of main **Figure 2C** experiment. (**C**) Quantitation of PPP1R15A mRNA levels in response to indicated stressors as determined by real-time qPCR. (**D**) RNA-seq data analysis. Smear plot in which top 40 differentially expressed genes are highlighted in red (control versus poly(I:C) transfection for 16 hrs). For clarity, all cell lines used in this study are combined in this plot. The position of PPP1R15A (not within top 40) is also shown as a reference (brown). (**E**) Splicing analysis shows no aberrant splicing patterns in the mutant cell lines. For clarity, reads mapped to standard reference genome (gray) and mutation specific references (colored) are shown separately (only pertinent region is shown). The distance between the end of uORF and donor-splice site is 17 bp. (**F**) Western blot for PPP1R15A and eIF2α under indicated conditions. (**G**) Polyribosome profiling of short uORF cell line versus wild-type 16 hrs after poly(I:C) transfection. (**H**) Proposed model of changes in phosphorylated eIF2α, PPP1R15A, and overall translation under indicated conditions for mild, modest, and severe stress.

**Supplemental Figure 4**

(**A**) Mouse body weight. (**B**) H&E histology of various organs is shown for Ppp1r15a uORF mutant and wild-type mice.

**Supplemental Figure 5**

(**A**) Polyribosome profiling of brain extracts from mice treated with 5 mg/kg LPS i.v. for indicated durations. We note that the resolution of mouse brain polyribosome profiling is suboptimal, consistent with a report by others.^73^ (**B**) Smear plot for Ppp1r15a uORF mutant versus wild-type. Top 20 differentially expressed genes are highlighted in red (ribo-seq data). (**C**) Select cytokine/chemokine levels are shown (ribo-seq data; the list is based on KEGG pathway mmu04060 and lowly expressed genes are filtered). (**D**) Heatmap of select antiviral genes under indicated conditions as determined by RNA-seq. (**E**) Polyribosome profiling of liver is shown.

**Supplemental Figure 6**

(**A**) Schematic of Ribo-seq workflow. (**B**) Genome-wide codon coverage analysis shows no overt differences in ribosome occupancy of codons among the conditions. (**C**) Histogram of ribosome footprint positions relative to coding sequences genome-wide is shown. The preponderance of red color indicates enrichment of frame specific ribosome footprints.

**Supplemental Figure 7**

(**A**) The effect of ASO2 on PPP1R15A protein levels is shown. uORF mutant cells (No uORF clone) and wild-type 293T with scramble ASO were used as positive and negative controls, respectively. Thapsigargin was added to induce PPP1R15A mRNA. (**B**) Western blot analysis of Ppp1r15a. Mice were injected with 5 mg/kg LPS i.v. followed by 10 mg/kg ASO i.v. at indicated time points and tissues were harvested 24 hrs after LPS. ASO2 scramble is shown. (**C**) ASO intervention at 14 hrs after 5 mg/kg LPS iv.

**Supplemental Figure 8**

(**A**) Motif prediction analysis of Ppp1r15a uORF3 microprotein using Eukaryotic Linear Motif resource. (**B**) Schematic of experiments in which Twin-Strep-tag was pulled down using the Strep-Tactin system. Human PPP1R15A uORF2 corresponds to mouse uORF3. (**C**) Workflow used for PPP1R15A mRNA pulldown followed by mass spectrometry. Top 12 proteins detected are listed. (**D**) uORF peptides (1.5 μM) were transfected into indicated cell lines using Chariot Protein Delivery Reagent with or without poly(I:C) transfection for 2 hrs. (**E**) PPP1R15A protein levels were determined by western blot with or without uORF peptides in the culture medium (15 μM for 2 hrs). PP(+), PP(-) denote polyproline(+) and polyproline(-) cell lines, respectively. (**F**) Computational prediction of Ppp1r15a uORF3 microprotein structure. Two ab initio protein structure tools, Robetta (David Baker lab)^40^ and Quark (Yang Zhang lab)^41^, both resulted in the same alpha-helix structure. (**G**) Docking prediction of the uORF and PPP1R15A:Protein phosphatase 1 (PP1) complex using ClusPro.^42^ uORF microprotein is shown in yellow, PP1 in blue, and PPP1R15A (the C-terminus domain only) in turquoise. The crystal structure of PPP1R15A:PP1 complex was obtained from the Protein Data Bank (4XPN).^11^ The catalytic domain is highlighted in red (arrow). The uORF microprotein is predicted to block the entry groove to the PP1 catalytic domain (arrow). The uORF microprotein also makes a close contact with PPP1R15A (circle). (**H**) Immunoprecipitation of PP1 followed by mass spectrometry under indicated conditions.

**Supplemental Figure 9**

(**A**) Synthesized Ppp1r15a uORF peptide is shown. (**B**) Schematic of phage display method. (**C**) Sequencing results of enriched M13 bacteriophage. Docking simulation between uORF microprotein and peptides identified by phage display. Results for Top 2 lowest ΔG are shown.

**Supplemental Figure 10**

(**A**) MHC peptidomics workflow is shown. (**B**) Overlay of non-metric multidimensional scaling plots (NMDS) for MHC-I bound peptides derived from WT and Ppp1r15a uORF^-/-^ mouse kidneys per timepoint. (**C**) Venn diagrams showing the degree of MHC-I peptide overlap per indicated conditions. Only non-redundant, 9-mer peptides that were detected in all triplicates per condition with H2-Kb binding %rank below 2% are shown. Gene names for peptides that were detected in WT and Ppp1r15a uORF^-/-^ at all timepoints but did not overlap between the two genotypes are listed. Heatmaps represent enriched pathways based on 9-mer MHC-I peptides detected in all conditions per genotype.

**Supplemental Figure 11**

(**A** - **B**) Polr1a uORF translation as determined by Ribo-seq after *in vivo* lactimidomycin treatment. Lactimidomycin treatment enables identification of translation initiation sites because the compound blocks elongation (but not initiation), hence resulting in accumulation of initiating ribosomes at putative translation initiation sites. Reads mapped to the Polr1a transcript (ENSEMBL Polr1a-201) are shown. (**C** – **D**) Comparison of Ribo-seq data between wild-type and Ppp1r15a uORF^-/-^ mice, 16 hrs after LPS (without lactimidomycin). Note the prominence of red-color frame reads in WT corresponding to the Polr1a uORF but not in Ppp1r15a uORF^-/-^ mice.

