## Supplemental figures for "Translation rescue by targeting Ppp1r15a upstream open reading frame *in vivo*"

Suppl Fig 1

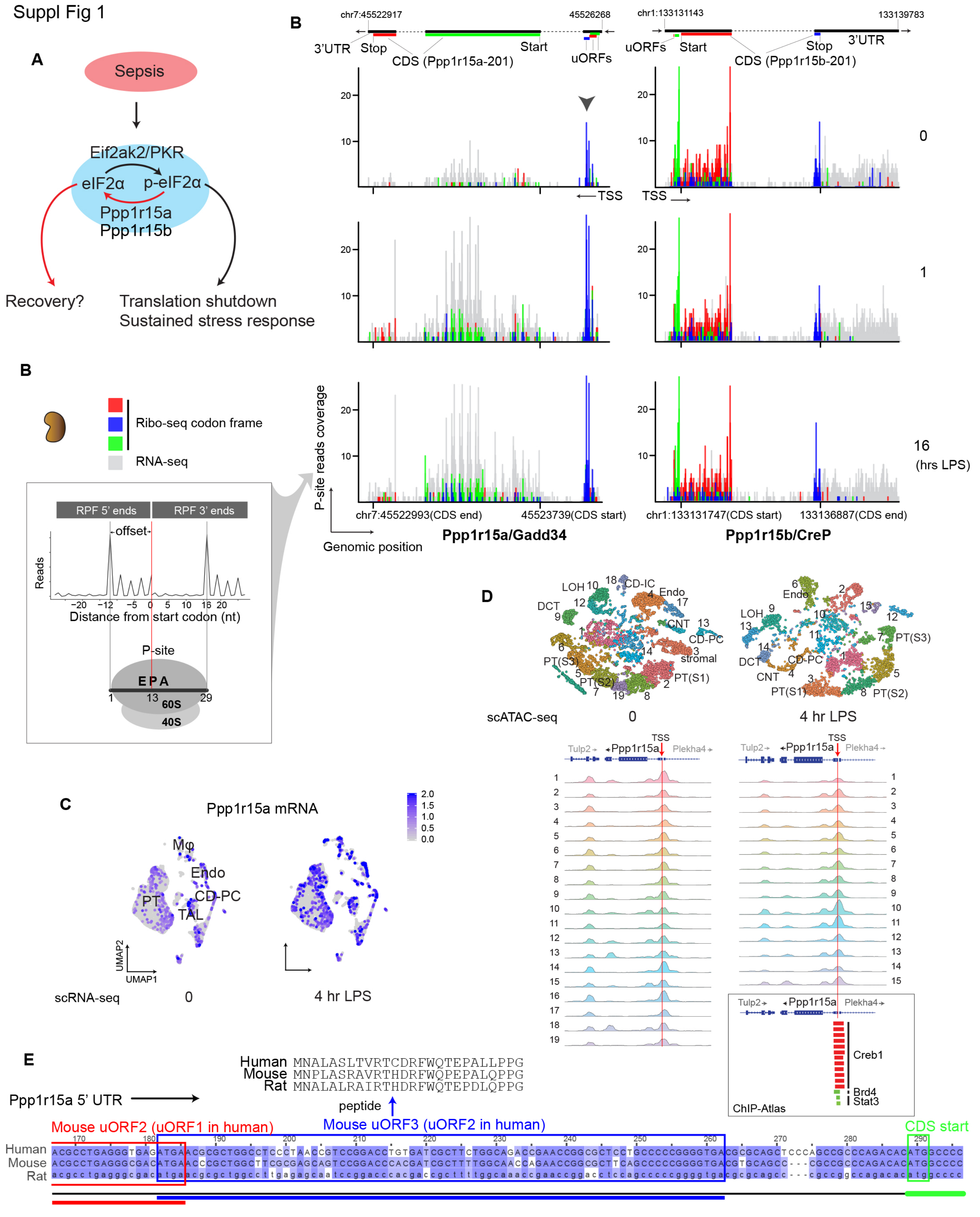


**Supplemental Figure 1.** (**A**) Schematic of the eIF2 axis. Only pertinent kinase (Eif2ak2/PKR) is shown for clarity. (**B**) Mouse kidney Ribo-seq data analysis. P-site offset was computed from ribosome-protected mRNA fragments mapped to transcriptome genome-wide (inset). P-site reads coverage is shown for Ppp1r15a and Ppp1r15b, and these reads are color coded based on their codon frame use. The codon-frame periodicity was calculated from the transcription start site (TSS) in the genome coordinate. Thus, codon frame colors can be different before and after an intronic region for a given coding sequence (CDS). Regular RNA-seq reads are color coded in gray. Note the low translation efficiency of Ppp1r15a (but not Ppp1r15b) as determined by the low ribo-seq-to-RNA-seq ratio over the CDS. Arrowhead in the left upper panel points to distinct 3-nulceotide periodicity throughout the 26 amino acid codons of the 3rd Ppp1r15a uORF (blue). (reanalysis of published data[12]) (**C**) Single-cell RNA-sequencing of mouse kidneys showing transcriptional increases of Ppp1r15a in a wide range of kidney cell types after LPS challenge (reanalysis of published data[13]). (**D**) Single-nuclear ATAC-sequencing (Assay for Transposase-Accessible Chromatin) of mouse kidneys demonstrating the accessibility of Ppp1r15a promoter region in all cell types at baseline and after LPS challenge (red vertical lines). Inset is a summary of transcription factors identified by ChIP-seq at the Ppp1r15a promoter region (ChIP-Atlas, filtered for mouse kidneys and the significance threshold is set at 200). (**E**) Human, mouse and rat Ppp1r15a transcripts are aligned and annotated.

Supplemental Figure 2

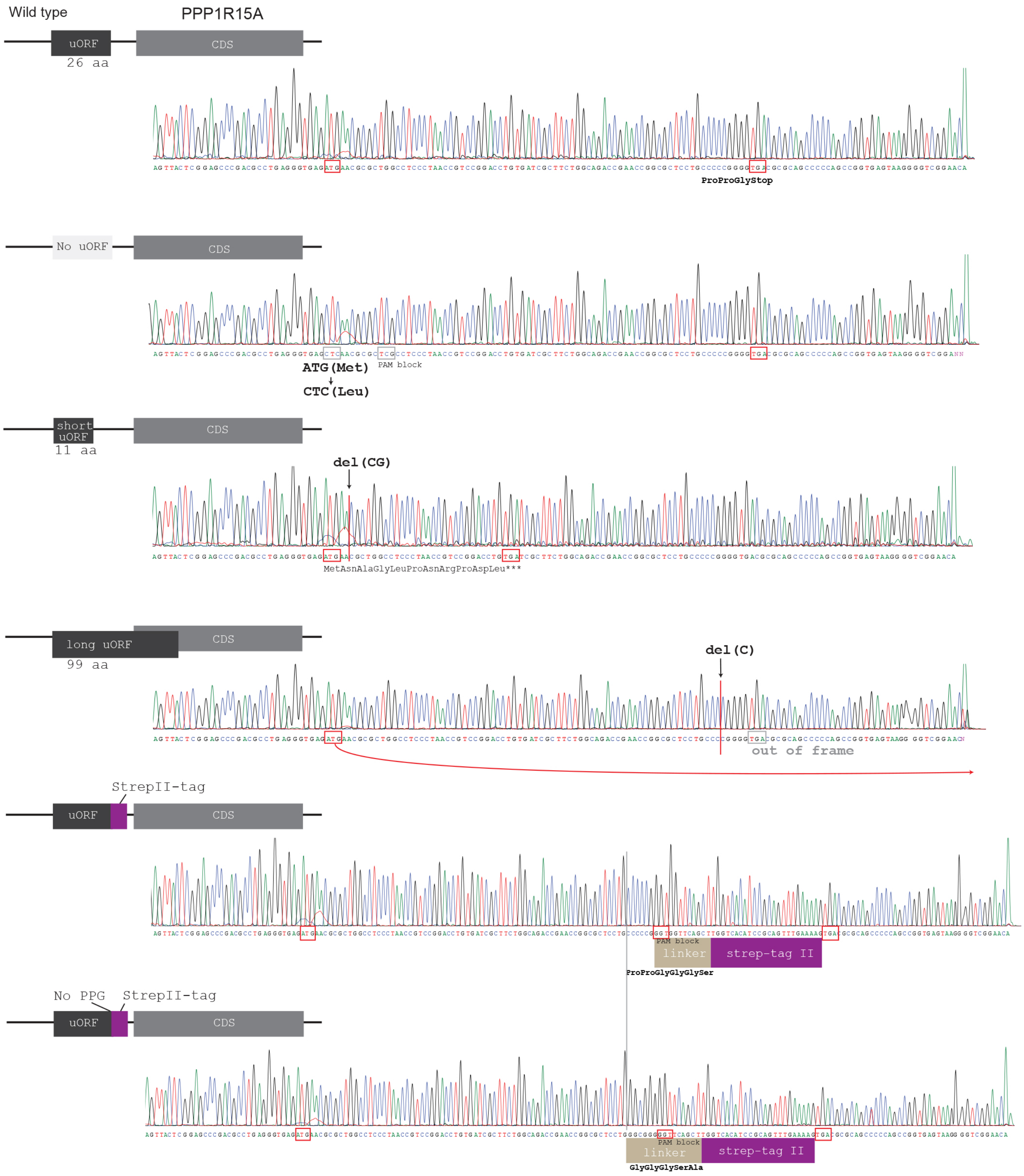


**Supplemental Figure 2**
Sanger sequencing chromatograms for all cell lines used in this study. Key features and mutations introduced by CRISPR/Cas9 are annotated.

Supplemental Figure 3

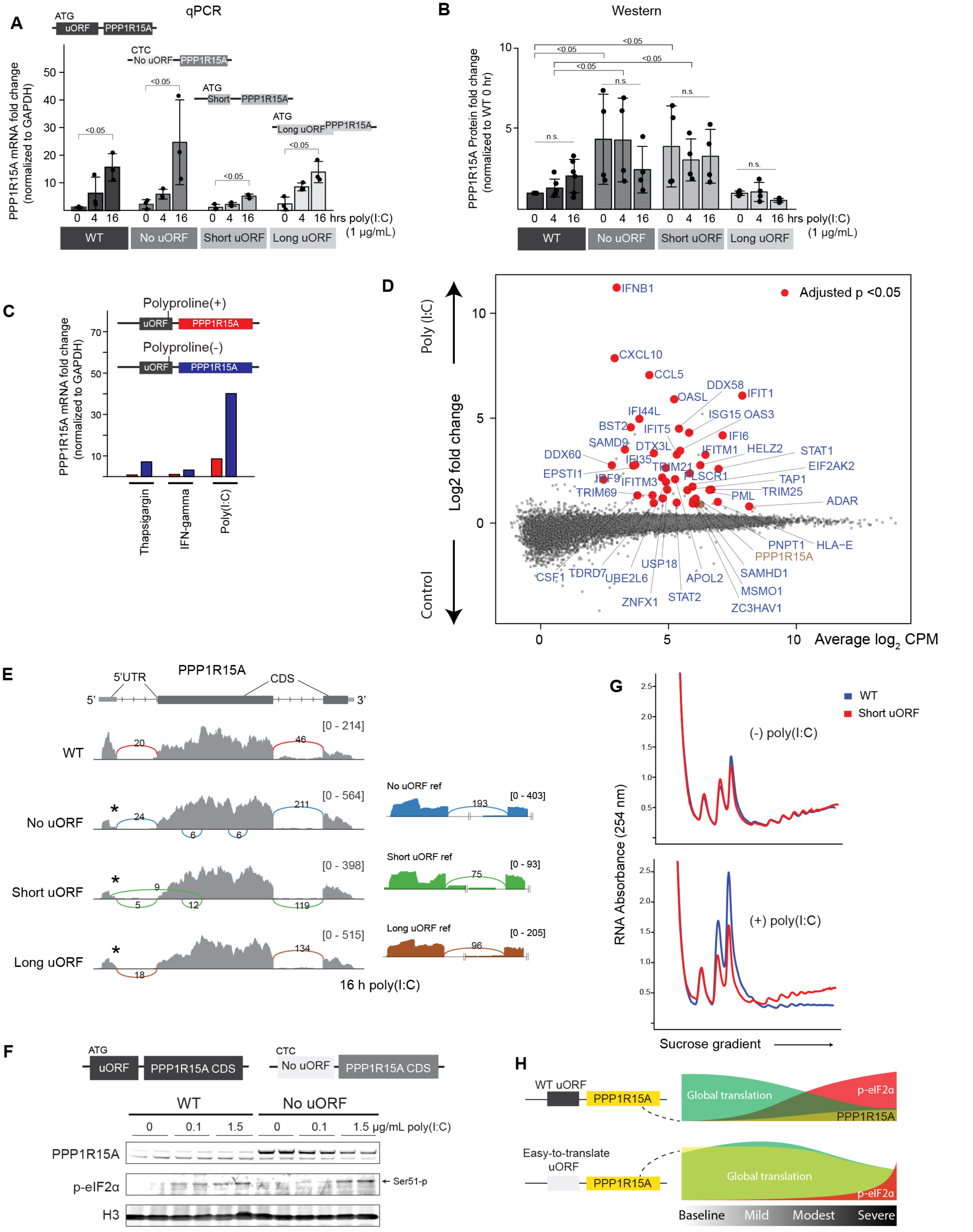


**Supplemental Figure 3.** (**A**) Quantitation of PPP1R15A mRNA levels as determined by real-time qPCR under indicated conditions. PCR primers used are identical to the one shown in main Figure 2C. n=3. (**B**) Quantitation of PPP1R15A protein levels as determined by western blot under indicated conditions. n=3, independent replicates of main Figure 2C experiment. (**C**) Quantitation of PPP1R15A mRNA levels in response to indicated stressors as determined by real-time qPCR. (**D**) RNA-seq data analysis. Smear plot in which top 40 differentially expressed genes are highlighted in red (control versus poly(I:C) transfection for 16 hrs). For clarity, all cell lines used in this study are combined in this plot. The position of PPP1R15A (not within top 40) is also shown as a reference (brown). (**E**) Splicing analysis shows no aberrant splicing patterns in the mutant cell lines. For clarity, reads mapped to standard reference genome (gray) and mutation specific references (colored) are shown separately (only pertinent region is shown). The distance between the end of uORF and donor-splice site is 17 bp. (**F**) Western blot for PPP1R15A and eIF2α under indicated conditions. (**G**) Polyribosome profiling of short uORF cell line versus wild-type 16 hrs after poly(I:C) transfection. (**H**) Proposed model of changes in phosphorylated eIF2α, PPP1R15A, and overall translation under indicated conditions for mild, modest, and severe stress.

Supplemental Figure 4

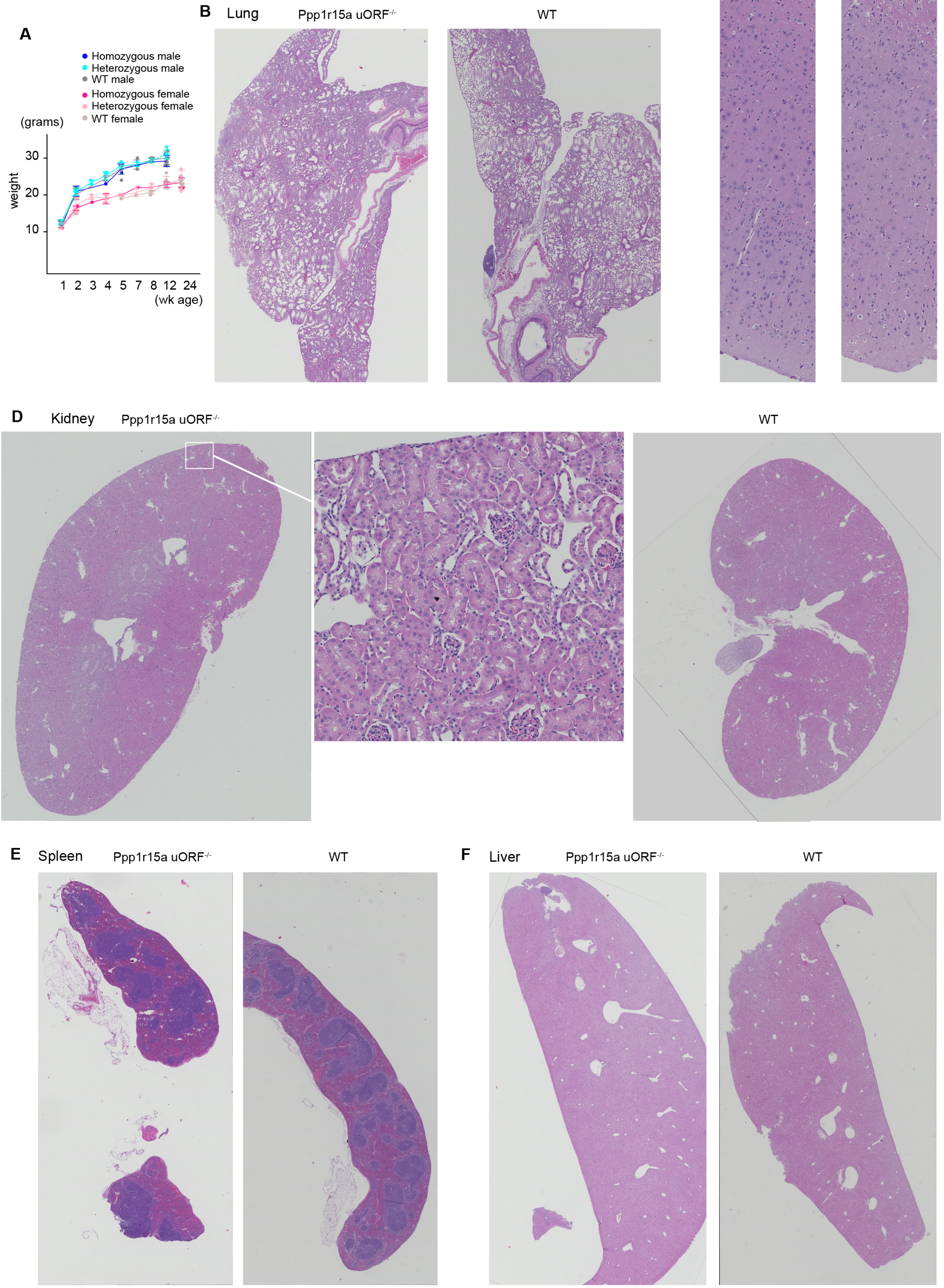


**Supplemental Figure 4.** (**A**) Mouse body weight. (**B**) H&E histology of various organs is shown for Ppp1r15a uORF mutant and wild-type mice.

Supplemental Figure 5

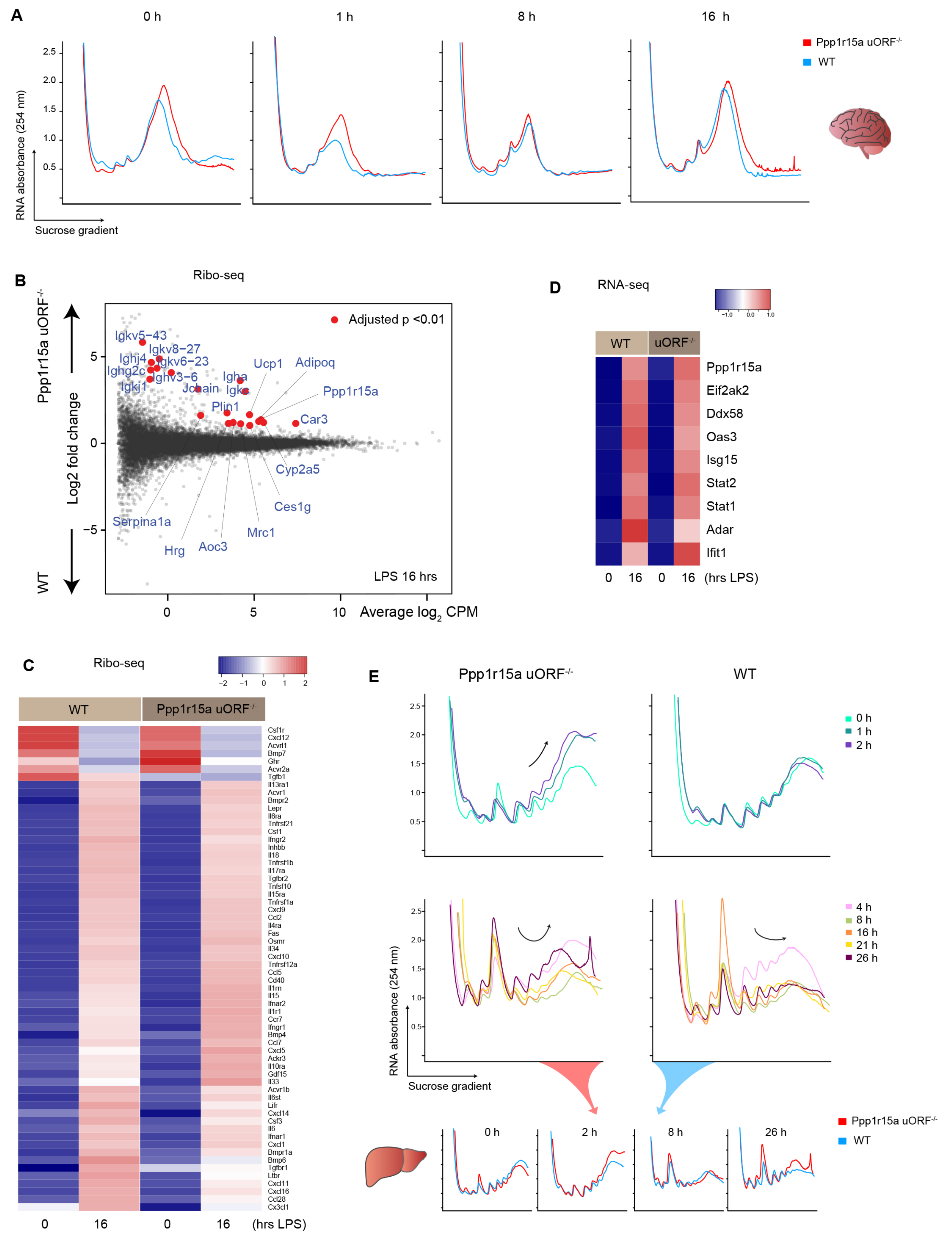


**Supplemental Figure 5.** (**A**) Polyribosome profiling of brain extracts from mice treated with 5 mg/kg LPS i.v. for indicated durations. We note that the resolution of mouse brain polyribosome profiling is suboptimal, consistent with a report by others.[73] (**B**) Smear plot for Ppp1r15a uORF mutant versus wild-type. Top 20 differentially expressed genes are highlighted in red (ribo-seq data). (**C**) Select cytokine/chemokine levels are shown (ribo-seq data; the list is based on KEGG pathway mmu04060 and lowly expressed genes are filtered). (**D**) Heatmap of select antiviral genes under indicated conditions as determined by RNA-seq. (**E**) Polyribosome profiling of liver is shown.

Supplemental Figure 6

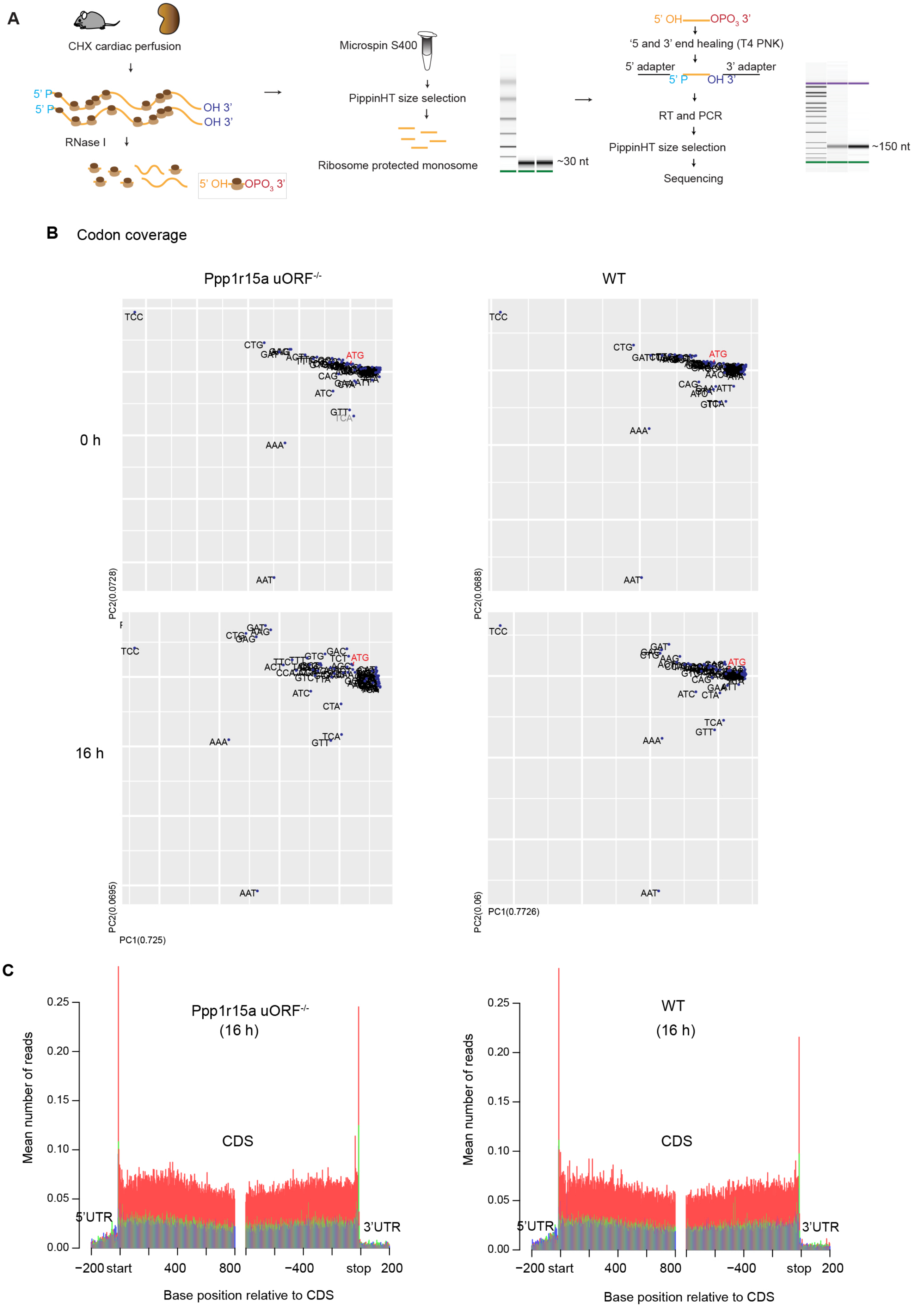


**Supplemental Figure 6.** (**A**) Schematic of Ribo-seq workflow. (**B**) Genome-wide codon coverage analysis shows no overt differences in ribosome occupancy of codons among the conditions. (**C**) Histogram of ribosome footprint positions relative to coding sequences genome-wide is shown. The preponderance of red color indicates enrichment of frame specific ribosome footprints.

Supplemental Figure 7

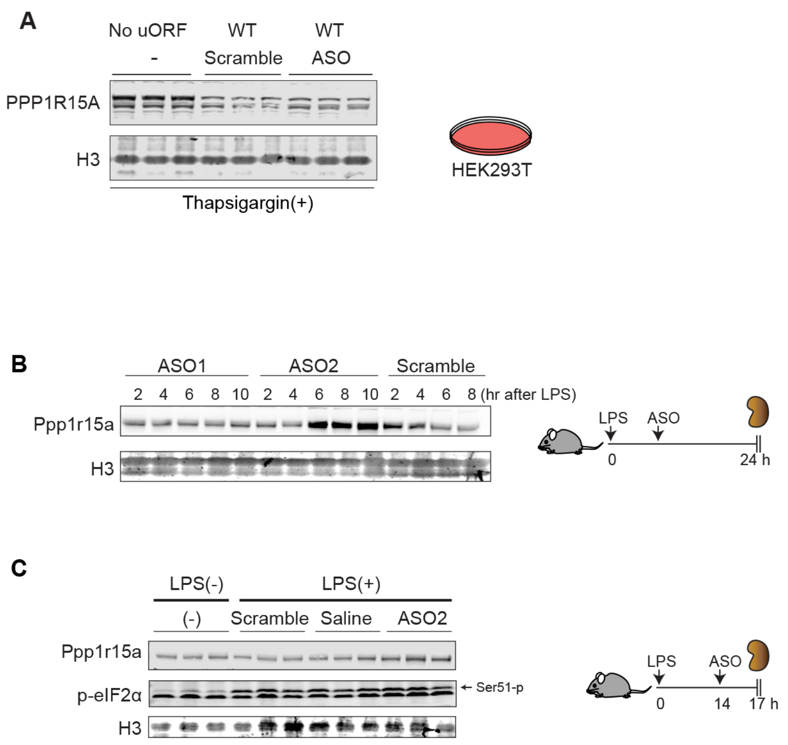


**Supplemental Figure 7**
(**A**) The effect of ASO2 on PPP1R15A protein levels is shown. uORF mutant cells (No uORF clone) and wild-type 293T with scramble ASO were used as positive and negative controls, respectively. Thapsigargin was added to induce PPP1R15A mRNA. (**B**) Western blot analysis of Ppp1r15a. Mice were injected with 5 mg/kg LPS i.v. followed by 10 mg/kg ASO i.v. at indicated time points and tissues were harvested 24 hrs after LPS. ASO2 scramble is shown. (**C**) ASO intervention at 14 hrs after 5 mg/kg LPS iv.

Supplemental Figure 8

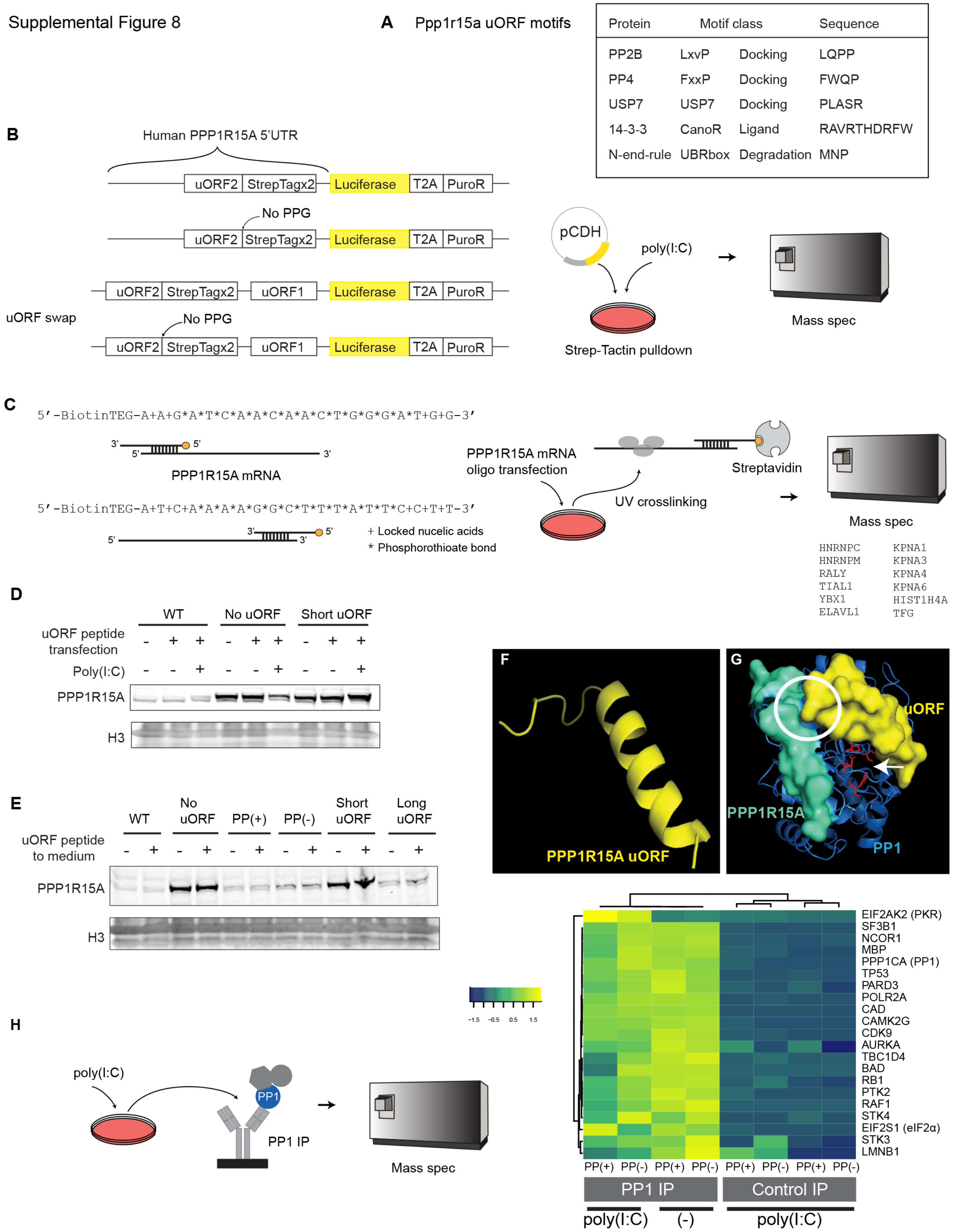


**Supplemental Figure 8.** (**A**) Motif prediction analysis of Ppp1r15a uORF3 microprotein using Eukaryotic Linear Motif resource. (**B**) Schematic of experiments in which Twin-Strep-tag was pulled down using the Strep-Tactin system. Human PPP1R15A uORF2 corresponds to mouse uORF3. (**C**) Workflow used for PPP1R15A mRNA pulldown followed by mass spectrometry. Top 12 proteins detected are listed. (**D**) uORF peptides (1.5 μM) were transfected into indicated cell lines using Chariot Protein Delivery Reagent with or without poly(I:C) transfection for 2 hrs. (**E**) PPP1R15A protein levels were determined by western blot with or without uORF peptides in the culture medium (15 μM for 2 hrs). PP(+), PP(-) denote polyproline(+) and polyproline(-) cell lines, respectively. (**F**) Computational prediction of Ppp1r15a uORF3 microprotein structure. Two ab initio protein structure tools, Robetta (David Baker lab)[40] and Quark (Yang Zhang lab)[41], both resulted in the same alpha-helix structure. (**G**) Docking prediction of the uORF and PPP1R15A:Protein phosphatase 1 (PP1) complex using ClusPro.[42] uORF microprotein is shown in yellow, PP1 in blue, and PPP1R15A (the C-terminus domain only) in turquoise. The crystal structure of PPP1R15A:PP1 complex was obtained from the Protein Data Bank (4XPN).[11] The catalytic domain is highlighted in red (arrow). The uORF microprotein is predicted to block the entry groove to the PP1 catalytic domain (arrow). The uORF microprotein also makes a close contact with PPP1R15A (circle). (**H**) Immunoprecipitation of PP1 followed by mass spectrometry under indicated conditions.

Supplemental Figure 9

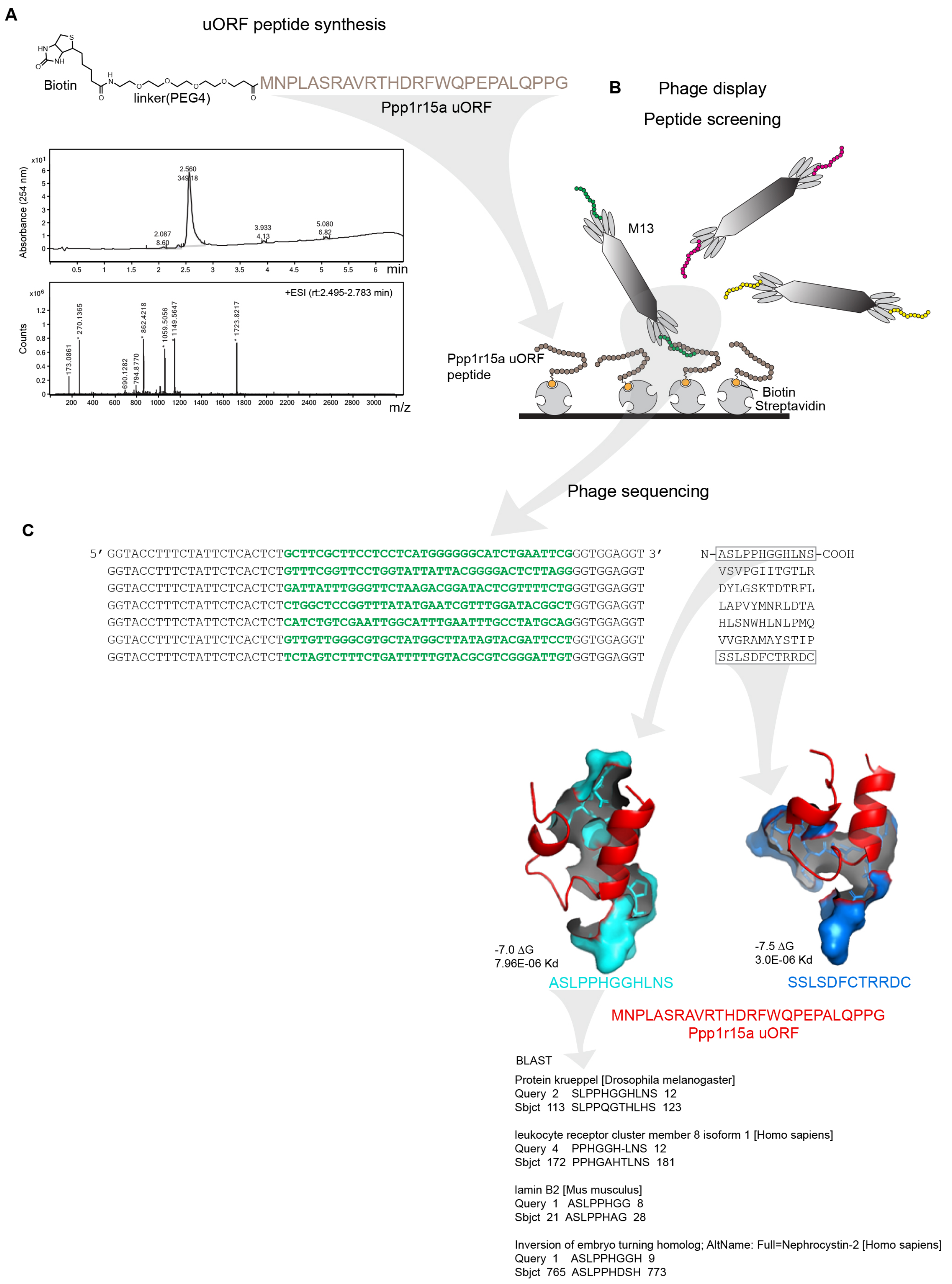


**Supplemental Figure 9**
(**A**) Synthesized Ppp1r15a uORF peptide is shown. (**B**) Schematic of phage display method. (**C**) Sequencing results of enriched M13 bacteriophage. Docking simulation between uORF microprotein and peptides identified by phage display. Results for Top 2 lowest ΔG are shown.

Supplemental Figure 10

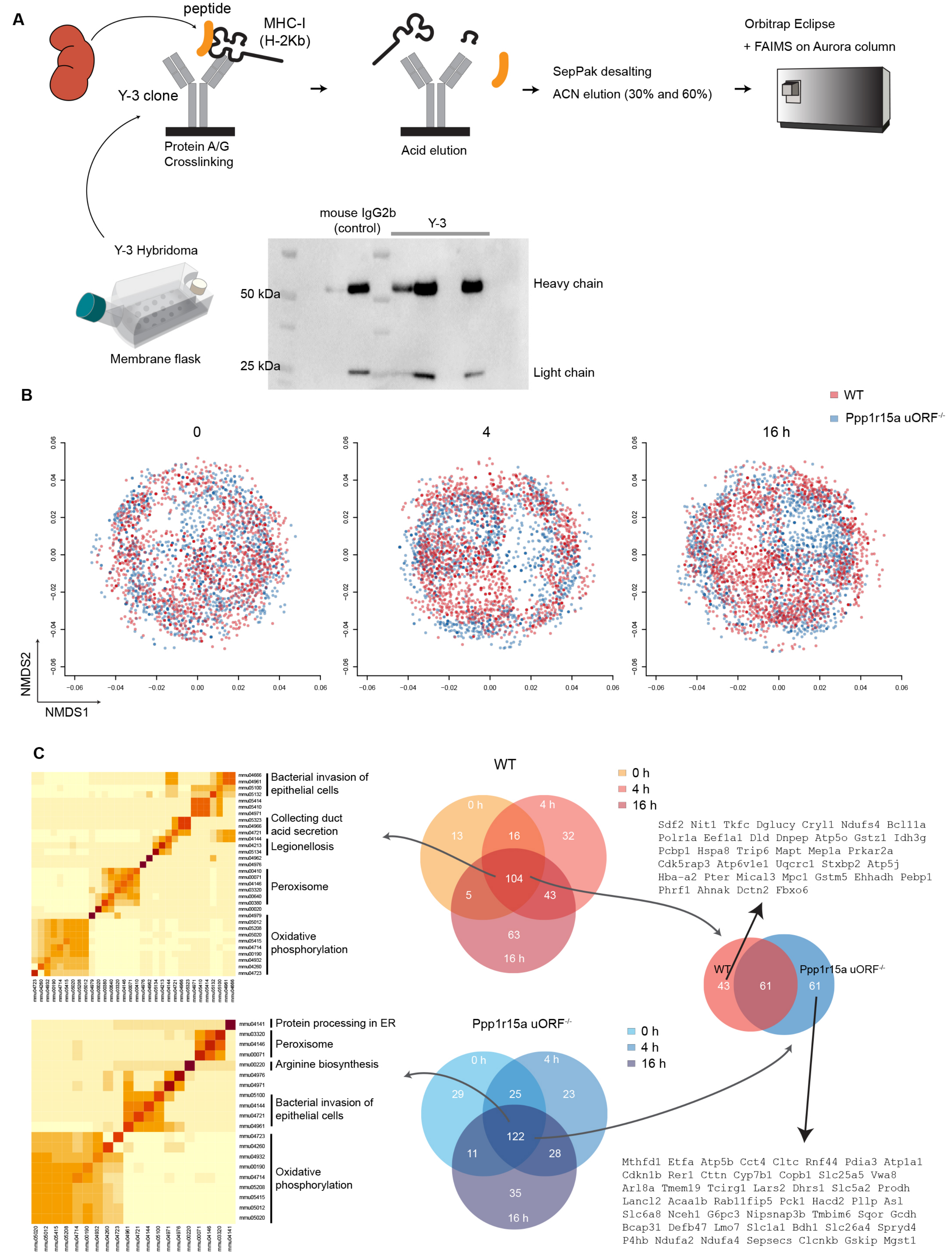


**Supplemental Figure 10.** (**A**) MHC peptidomics workflow is shown. (**B**) Overlay of non-metric multidimensional scaling plots (NMDS) for MHC-I bound peptides derived from WT and Ppp1r15a uORF-/- mouse kidneys per timepoint. (**C**) Venn diagrams showing the degree of MHC-I peptide overlap per indicated conditions. Only non-redundant, 9-mer peptides that were detected in all triplicates per condition with H2-Kb binding %rank below 2% are shown. Gene names for peptides that were detected in WT and Ppp1r15a uORF-/- at all timepoints but did not overlap between the two genotypes are listed. Heatmaps represent enriched pathways based on 9-mer MHC-I peptides detected in all conditions per genotype.

Supplemental Figure 11

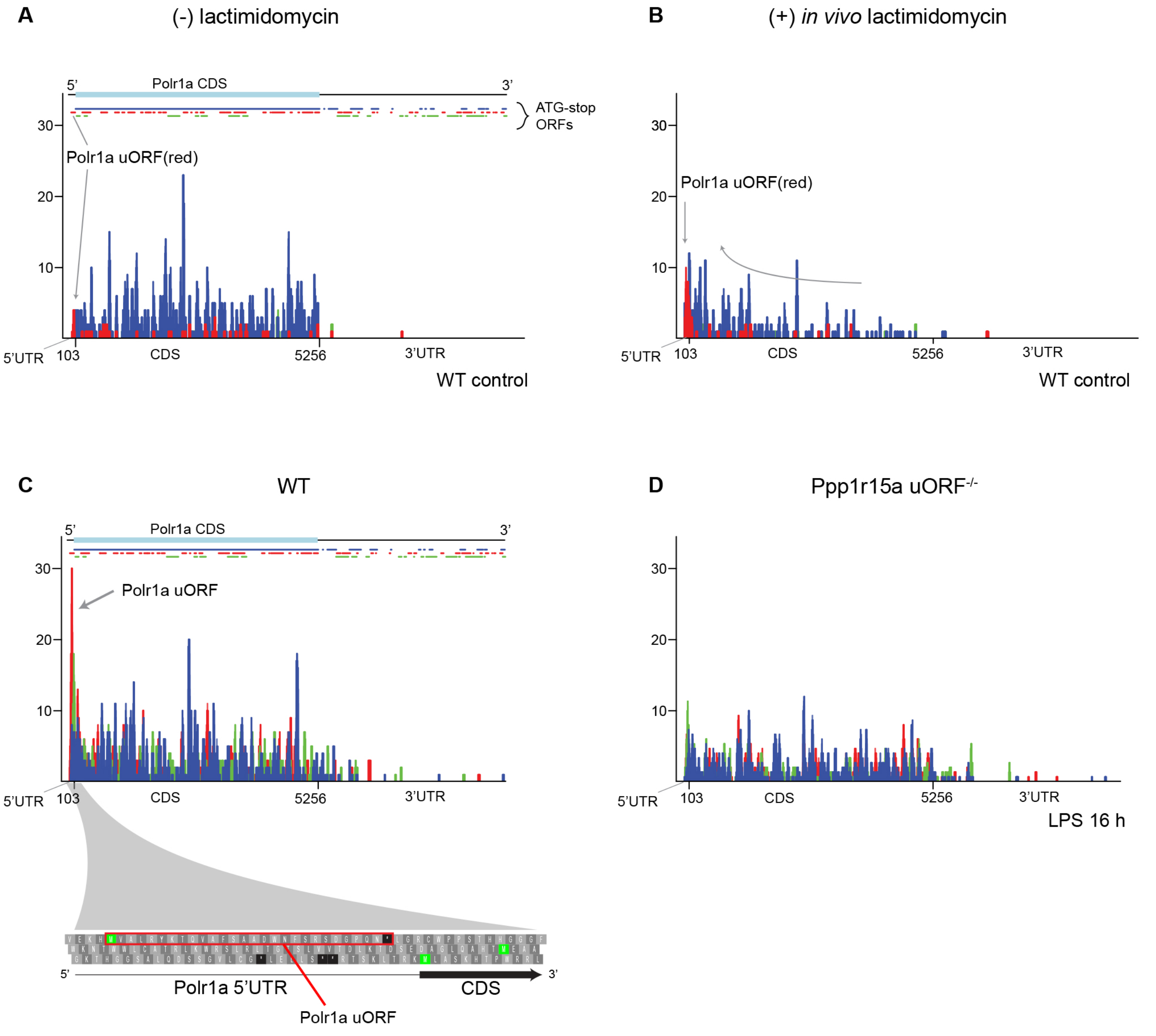


**Supplemental Figure 11**
(**A** - **B**) Polr1a uORF translation as determined by Ribo-seq after in vivo lactimidomycin treatment. Lactimidomycin treatment enables identification of translation initiation sites because the compound blocks elongation (but not initiation), hence resulting in accumulation of initiating ribosomes at putative translation initiation sites. Reads mapped to the Polr1a transcript (ENSEMBL Polr1a-201) are shown. (**C** – **D**) Comparison of Ribo-seq data between wild-type and Ppp1r15a uORF-/- mice, 16 hrs after LPS (without lactimidomycin). Note the prominence of red-color frame reads in WT corresponding to the Polr1a uORF but not in Ppp1r15a uORF-/- mice.
